# The mechanism of nicotinamide phosphoribosyltransferase whereby positive allosteric modulation elevates cellular NAD^+^

**DOI:** 10.1101/2022.10.21.513220

**Authors:** Kiira M. Ratia, Zhengnan Shen, Jesse Gordon-Blake, Hyun Lee, Megan S. Laham, Isabella S. Krider, Nicholas Christie, Martha Ackerman-Berrier, Christopher Penton, Natalie G. Knowles, Soumya Reddy Musku, Jiqiang Fu, Ganga Reddy Velma, Rui Xiong, Gregory R J Thatcher

**Affiliations:** Department of Pharmaceutical Sciences, College of Pharmacy, University of Illinois at Chicago (UIC), Chicago, IL 60612, USA; Research Resources Center, University of Illinois at Chicago (UIC), Chicago, IL 60612, USA; Department of Pharmacology & Toxicology, College of Pharmacy, University of Arizona, Tucson, AZ 85721, USA; Department of Chemistry & Biochemistry, Colleges of Science and Medicine, University of Arizona, Tucson, AZ 85721, USA

## Abstract

In aging and disease, cellular NAD^+^ is depleted by catabolism to nicotinamide (NAM) and NAD^+^ supplementation is being pursued to enhance human healthspan and lifespan. Activation of nicoti namide phosphoribosyl -transferase (NAMPT), the rate-limiting step in NAD^+^ biosynthesis, has potential to increase salvage of NAM. Novel NAMPT positive allosteric modulators (N-PAMs) were discovered in addition to demonstration of NAMPT activation by biogenic phenols. The mechanism of activation was revealed through synthesis of novel chemical probes, new NAMPT co-crystal structures, and enzyme kinetics. Binding to a rear channel in NAMPT regulates NAM binding and turnover, with biochemical observations being replicated by NAD^+^ measurements in human cells. The mechanism of action of N-PAMs identifies, for the first time, the role of the rear channel in regulation of NAMPT turnover coupled to feedback inhibition by NAM. N-PAM inhibition of low affinity, non-productive NAM binding via the rear channel, causes a right-shif t in K_I_(NAM) that accompanies an increase in enzyme activity. Conversion of an N-PAM to a high-affinity ligand blocks both high and low affinity NAM binding, ablating enzyme activity. In the presence of an N-PAM, NAMPT boosts NAD^+^ biosynthesis at higher NAM concentrations, in addition to relieving inhibition by NAD^+^. Since cellular stress often leads to enhanced catabolism of NAD^+^ to NAM, this mechanism is relevant to supporting cellular N AD^+^ levels in aging and disease. The tight regulation of cellular NAMPT is differentially regulated by N-PAMs and other activators, indicating that different classes of pharmacological activators may be engineered for cell and tissue selectivity.

## INTRODUCTION

Nicotinamide adenine dinucleotide (NAD^+^) and its 2-phosphate derivative (NADP^+^) are formed on electron transfer from NADH and NADPH, respectively. The electron transfer reactions regulated by these enzyme cofactors drive cellular metabolic processes. NAD^+^ serves as a substrate for important enzymes that catabolize NAD^+^ to nicotinamide (NAM), a process that can lead to severe disruption of the cellular NAD^+^ economy. This disruption and depletion of NAD^+^ is closely associated with aging and metabolic disorders; and therefore, replenishment of cellular NAD^+^ by “NAD^+^-enhancing drugs” has evolved as a strategy for therapeutics directed at enhancement of lifespan and healthspan (Garten et al., 2015; J. Yoshino, Baur, & Imai, 2018). NAD^+^ supplementation as an antiaging therapy has been reported in the popular press, accompanied by contemporary clinical trials on NAD^+^ dietary supplements (Dellinger et al., 2017; Martens et al., 2018). Administration of NAM or nicotinamide mononucleotide (NMN), was shown to improve aspects of healthspan or to increase lifespan in mice (Mitchell et al., 2018; Uddin, Youngson, Doyle, Sinclair, & Morris, 2017). Recently, treatment of prediabetic postmenopausal women for 10 weeks with NMN was shown to restore muscle insulin sensitivity and insulin signaling (M. Yoshino et al., 2021). NMN is a biosynthetic precursor of NAD^+^ and is the product of the reaction of NAM with α-D-5-phosphoribosyl-1-pyrophosphate (PRPP) catalyzed by the enzyme nicotinamide phosphoribosyltransferase (NAMPT) (Garten et al., 2015).

Accumulated DNA damage associated with normal aging leads to elevated poly-ADP-ribose polymerase (PARP) activity: PARP consumes NAD^+^ to ADP-ribosylate proteins at sites of DNA damage (Fang et al., 2014). A second source of NAD^+^ catabolism to NAM is NAD nucleosidase activity associated with CD38, the expression of which increases in multiple tissues with age and is associated with chronic inflammation associated with aging (inflammaging) and cellular senescence (Camacho-Pereira et al., 2016; Chini et al., 2020; Imai & Guarente, 2014; J. Yoshino et al., 2018). In response to the cellular stress induced by PARPs and CD38, upregulation of sirtuins (SIRTs) can provide cellular protection; however, SIRTs themselves catabolize NAD^+^ in the process of protein de-acylation. NAD^+^ depletion leads to reduced SIRT1-mediated deacetylation of PGC-1α, which results in defective mitochondria and increased release of reactive oxygen species that cause further DNA damage (Guarente, 2014; Imai & Guarente, 2014).

Mitochondrial DNA damage leads to age-related mitochondrial dysfunction and SIRT1-dependent triggering of hypoxic programs under normoxic conditions, events that can be reversed by restoring NAD^+^ (Gomes et al., 2013). Other ADP-ribosyltransferases (ARTs) contributing to NAD^+^ depletion include sterile alpha and TIR motif containing 1 (SARM1) (Essuman et al., 2017). Quantitatively, both PARP activity and SIRT activity each account for one third of NAD^+^ consumption (Liu et al., 2018).

In mammals, the rate-limiting step in NAD^+^ biosynthesis is the salvage of NAM and conversion to NMN, catalyzed by NAMPT (**Figs 1A,B**). NAMPT activity is viewed as key in cellular defense mechanisms controlling cell survival and maintaining metabolic homeostasis. ^2^ Female *Nampt+/*- mice have glucose tolerance and impaired insulin secretion that is ameliorated by administration of NMN.(Revollo et al., 2007; Stromsdorfer et al., 2016; J. Yoshino, Mills, Yoon, & Imai, 2011) NAMPT mediates an adaptive response to inflammatory, oxidative, and genotoxic stress, a response that declines with age, along with a reported significant decline in NAMPT and NAD^+^ levels.(Parihar & Brewer, 2007; J. Yoshino et al., 2011; Zhu, Lu, Lee, Ugurbil, & Chen, 2015) Although the role of NAMPT in physiology and pathophysiology is complex, the possibility of addressing diseases of normal aging, such as Alzheimer’s disease and related dementia (ADRD) and diseases of accelerated aging, such as Type-2 diabetes (T2D), by enhancing NAMPT activity, is compelling.

**Figure 1.**
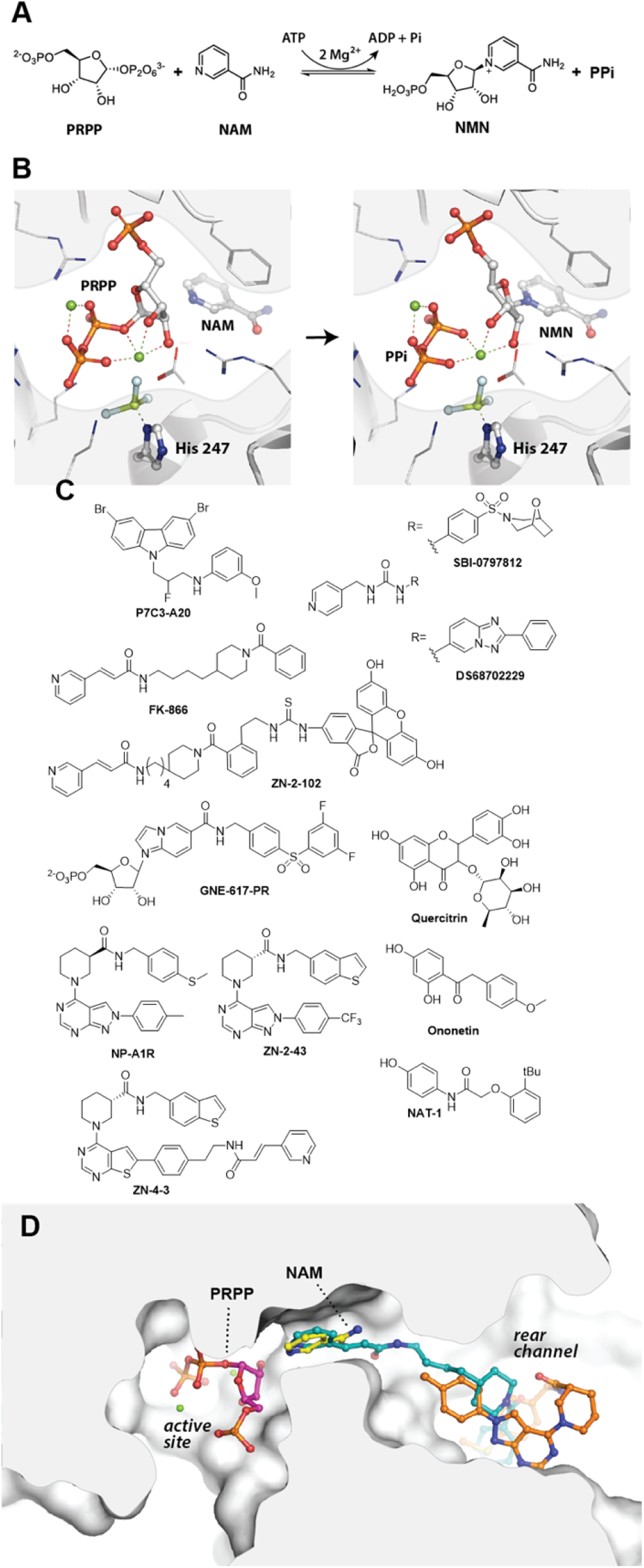
Enzyme and ligand structures. Conversion of NAM to NMN at the NAMPT active site requires His-247 phospho-nylation by ATP (A). Superposition of crystal structures illustrating the NAMPT active site (two Mg^2+^ ions shown as green dots) (PDB: 3DKL; 3DHF) (B). Structures of putative NAMPT activators, NAMPT inhibitors, and fluorescence-polarization (FP) probe ZN-2-102 (C). D. Superposition of FK866 (cyan) and NP-A1 (gold) bound to the rear channel of NAMPT (PDB: 2GVJ; 3DKL; 3DHF).

NAMPT catalyzes the formation of NMN and an ATPase reaction, both at the same active site, in a mechanism delineated in three papers from Schramm, Burgos and coworkers using: 1) kinetic analysis;(Burgos & Schramm, 2008) 2) structural biology;(Burgos, Ho, Almo, & Schramm, 2009) and 3) kinetic isotope effects (Burgos, Vetticatt, & Schramm, 2013). The chemical equilibrium modulated at the active site of NAMPT (PRPP + NAM ⇋NMN + PPi) lies to the left in the absence of ATP: i.e. slow conversion of NMN to NAM (**Figs 1A,B**). The ATPase reaction transferring the γ-phosphate of Mg^2+^-ATP to form *N*phosphohistidine (phospho-H247) occurs at the same active site, with nucleobases of ATP/ADP and NAM/NMN sequentially occupying a nucleobase binding pocket. The formation of phospho-H247 leads to a shift in the equilibrium to the right, resulting in a 1,100-fold increased efficiency for NMN formation.(Burgos et al., 2013) The increase in k_cat_/K_M_ is driven by a ≤ 170-fold enhancement of PRPP-dependent NAM binding and an increase in k_cat_. Importantly, phospho-H247 is required to coordinate two Mg^2+^ ions that stabilize the transition state for phosphoribosyl group transfer from the ribose anomeric carbon of PRPP to the pyridyl-*N* of NAM (Fig. 1B). Feedback inhibition is observed by both NAM and NAD^+^ and under most conditions, the ATPase reaction is not tightly coupled to turnover of NAM, leading to a seemingly inefficient catabolism of cellular ATP, which is amplified in the presence of NAM and the absence of PRPP. (Burgos & Schramm, 2008) A phenotypic screen of 1,000 compounds for neuronal cell viability yielded a carbazole hit that was subsequently claimed to be a NAMPT ligand and activator (P7C3-A20; Fig. 1C).(Pieper & McKnight, 2019) However, P7C3-A20 does not increase enzymic activity of recombinant NAMPT; as demonstrated in a recent report (Gardell et al., 2019) and by data reported herein. The 4-pyridyl compound, SBI-797812 (Fig. 1C),(Gardell et al., 2019) is an authentic NAMPT activator, from which further 4-pyridyl activators were derived, including DS68702229 that showed positive effects in an obesogenic mouse model. (Akiu et al., 2021) However, the 3-pyridyl analogue of SBI-797812 inhibits NAMPT(Zheng et al., 2013); indeed, 3-pyridyl and related NAMPT inhibitors are widely studied, since NAMPT is a target for inhibition in cancer therapy.(Chowdhry et al., 2019; Oh et al., 2014; Wilsbacher et al., 2017) Multiple co-crystal structures demonstrate the binding of potent NAMPT inhibitors in a “rear-channel” that accesses the nucleobase binding site from the opposite direction to the *N-*phosphohistidine-Mg^2+^ complexed PRPP, with most inhibitors (e.g. FK866) containing a nitrogenous base that binds in the nucleobase site, displacing NAM (Fig. 1D). Despite mechanistic studies on SBI-797812, the binding mode leading to NAMPT activation has not been fully defined.(Gardell et al., 2019)

In HTS assay of small molecule libraries, we identified more than one class of small molecule able to increase enzymic activity of recombinant NAMPT. (Gordon-Blake et al., 2019) Several phenolic compounds, including quercitrin and other biogenic and/or bioactive phenols were efficacious, with modest potency and binding affinity (**Fig. 1C**). These were differentiated from both: i) novel NAMPT positive allosteric modulators (N-PAMs) that bind NAMPT to increase activity in biochemical and cell-based assays; and from the SBI-class of activators (**Fig. 1C**). The N-PA Ms are also differentiated in their ability to couple the turnover of NAM → NMN with the ATPase activity of NAMPT. We were able to obtain multiple co-crystal structures of quercitrin and N-PAMs bound to the rear channel of NAMPT (**Fig. 1D**) and further able to co-crystalize a potent, novel NAMPT inhibitor by simple derivatization of the N-PAM. Structural analysis, combined with enzyme kinetics, support a novel mechanism of allosteric activation of NAMPT by regulation of productive and non-productive binding of NAM and mechanical coupling of the rear channel to the active site.

This work provides the structural and mechanistic basis for the design of small molecules as NAMPT activators of potential therapeutic value. It also defines a role for the rear channel that is unique to NAMPT among phosphoribosyltransferases, raising the question of possible endogenous activators and the contribution of NAMPT activation to the biological activity of biogenic phenols.

## RESULTS & DISCUSSION

### Activation of NAMPT

The primary assay of NAMPT enzyme activity measured NAD^+^ production by coupling the NAMPT-catalyzed production of NMN with: (i) the conversion of NMN to NAD^+^ catalyzed by nicotinamide/nicotinic acid mononucleotide adenylyltransferase (NMNAT); and (ii) alcohol dehydrogenase (ADH) and DT-diaphorase (**Fig 2A,B**). Optimized for HTS, the assay was used to screen: 20,000 compounds from ChemDiv libraries; and 2,000 compounds from the Microsource Spectrum library. The most active validated hit from the ChemDiv library was NP-A1 and validated hits from the Spectrum library included the flavonoid glycoside, quercitrin, onenetin, genestein, and dienestrol (**Fig. 2C; Fig. S1**). The aglycone, quercitin, was devoid of activity at ≤ 40 µM. Further examination of the isomers of NP-A1 revealed that the *R-*isomer was the most potent compound (EC_50_ = 37 nM), whereas the *S-*isomer gave the maximum observed activation of NAMPT relative to vehicle control (**Fig. 2C**). The trapping of NMN as a fluorescent adduct was the basis of an orthogonal assay for NAMPT activity, (Zhang et al., 2011) validated using the NAMPT inhibitor FK866 (**Fig. 2D**). This is an endpoint assay in which NMN accumulates, in contrast to the coupled NAMPT assay in which NMN is immediately captured by NMNAT. In neither NAMPT assay did PC73-A20 show activation of NAMPT (data not shown). After completion of our HTS campaign, the 4-pyridyl activator, SBI-0797812, was reported,(Gardell et al., 2019) which was compared with N-PAMs and quercitrin and shown to activate NAMPT (**Fig. 2E**).

**Figure 2.**
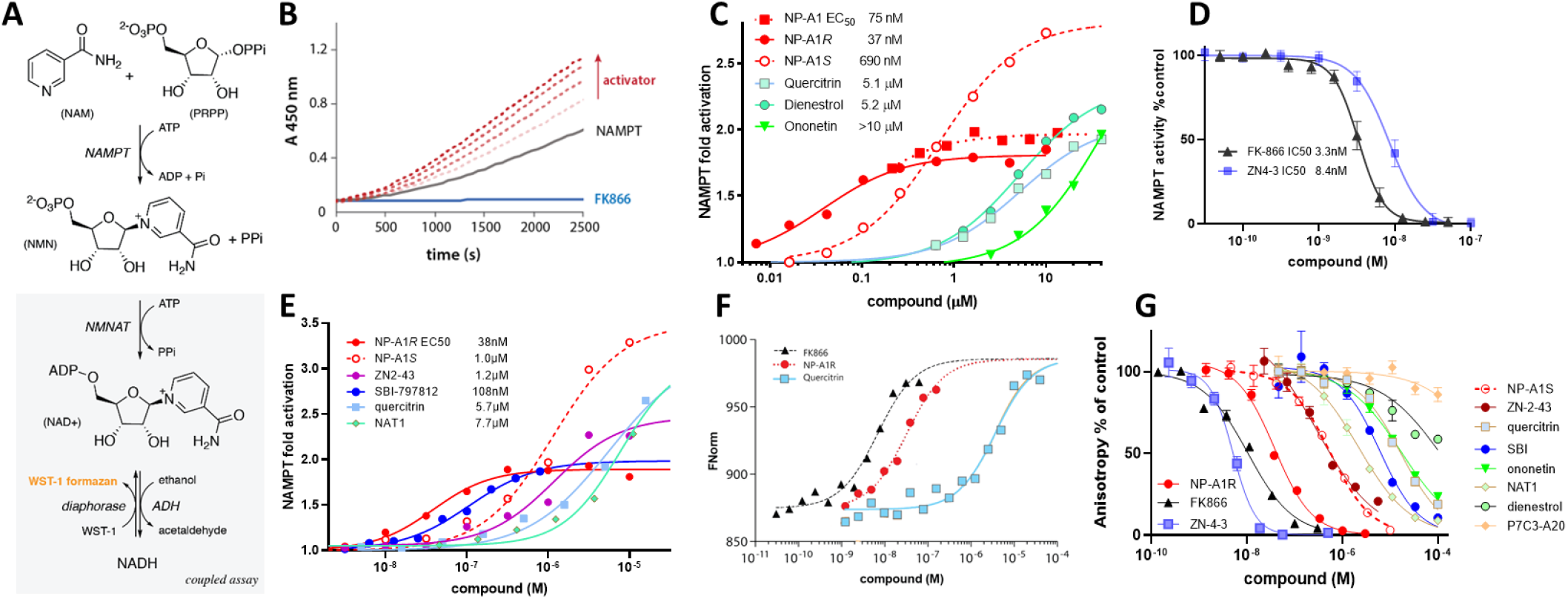
NAMPT activation and binding. Coupled primary enzyme assay used for HTS (A), with raw kinetic traces showing activ atio n (B), and derived concentration-response curves for HTS hits and NP-A1 isomers (C). Orthogonal NAMPT assay validated with NAMPT inhibitors (D) and derived concentration-response curves for activators and N-PAMs (E). Binding to NAMPT measured by MST (F) an d displacement of the rear channel FP-probe ZN-2-102 (G).

### Binding to NAMPT

Orthogonal biophysical assays were used to confirm and quantify ligand binding to NAMPT. Microscale thermophoresis (MST) measurements replicated the order of binding affinity indicated from NAMPT kinetic data: FK866 > NP-A1R > quercitrin (**Fig. 2F**). Given the occupation of the nucleobase binding site and rear channel by NAMPT inhibitors, it was straightforward to utilize an inhibitor scaffold to synthesize and validate a fluorescence polarization (FP) probe (ZN-2-102 **Fig. 1C**) to measure binding to NAMPT through displacement of the FP-probe (**Fig. 2G**). The combined kinetic and biophysical data demonstrated NP-A1R to be a high affinity ligand with modest maximal activation and NP-A1S to be a weaker ligand with superior maximal activation. Quercitrin, as representative of the phenolic activator class, was a weak binding NAMPT ligand with high activation at higher concentrations.

To confirm the binding to the NAMPT rear channel indicated by the FP-probe measurements and to understand the mechanism of enzyme activation by both N-PAMs and the bioactive phenolic activators represented by quercitrin, we obtained several co-crystal structures. In addition to structures with bound quercitrin, NP-A1R, and NP-A1S, we obtained structures with a synthetic N-PAM (ZN-2-43 **Fig. 1C**) and a novel inhibitor obtained from this N-PAM (ZN-4 3 **Fig. 1C**). The synthetic ligands were validated as either a NAMPT activator or NAMPT inhibitor in the orthogonal enzyme activity assay (**Figs 2D,E**). Importantly, all xenobiotic ligands occupy the same rear channel occupied by FK866, but activators do not extend into the nucleobase binding pocket (**Figs 1D, 3**).

### Structural analysis of NAMPT binding

Over 70 crystal structures of NAMPT are represented in the RCS-PDB database, the majority being structures of the human enzyme and with bound inhibitors. As shown in **Figure 1D**, 3-pyridyl inhibitors, such as FK866, bind in the “rear channel” with the pyridyl group occupying the nucleobase binding site and superimposing on NAM. Although displacement of NAM is sufficient for biochemical inhibition, it has been proposed that the most potent inhibitors also act as substrates undergoing phosphoribosylation (e.g. GNE-617-PR; **Fig. 1C**).(Oh et al., 2014) The nucleobase of both NA M and NAMPT inhibitors is bound in a Phe-193/Tyr-18 pi-pi clamp. The nucleobase pocket must also accommodate the adenine group during the ATPase reaction converting A TP to ADP: adenine binds with the nucleobase displaced and rotated approximately 100° with respect to the orientation of NAM using a Arg-311/Phe-193 cation-pi clamp. NAMPT functions as a homodimer in which the dimer interface provides two symmetrically mirrored active sites (**Fig. S2**). The rear channel, nucleobase pocket, and active site are largely lined by residues of the first monomer; whereas, the pi-pi clamp Tyr-18 and hydrophilic residues at the mouth of the active site are supplied by the second monomer. The rear channel of human NAMPT is formed by two small β-strands (β14 and β15) at the dimer interface. Despite strong structural and mechanistic similarities between NAMPT and human nicotinic acid phosphoribosyl-transferase (NAPRT), the rear channel of NAPRT, formed by β-strands β13 and β14 is occluded, blocking binding of FK866 and other NAMPT inhibitors. The rear channel of NAMPT is a seemingly unique feature.

NP-A1 structures were obtained with bound NAM and quercitrin structures contained either bound NAM or a stable ADP analogue (AMPCP). The structures of ZN - 2 - 43 and ZN-4-3 did not contain bound NAM. An unliganded NAMPT structure, with bound NAM, was also obtained for comparison. A key feature of the rear channel adjacent to the nucleobase binding site is a H-bonding network with three water molecules: two H-bonded to NAM, Ser-241, Ser-275, and Asp-219; and a third water molecule not directly H-bonded to any amino acid residue, which is clearly shown in the NP-A1S structure (**Fig. 3A)**. In the NAMPT structures without bound ligand, a fourth water molecule was observed, which was displaced by all ligands (**Fig. S3**). Superposition of FK866 with NAM and NP-A1S bound to the rear channel of NAMPT shows the displacement of two of the three water molecules, with the water interactions being replaced by H-bonding with the amide group of FK866 (**Fig. 3B**; an arrangement also observed in the structure of the novel NAMPT-bound inhibitor ZN-4-3) The phenolic 7-OH of quercitrin displaces one of the three water molecules in this network (**Fig. 3C**). In the two quercitrin co-crystal structures, the 7-hydroxy group sits 3. 5 Å and 4.1Å from the amide nitrogen of NAM and from the adenine ring nitrogen of ADP (AMPCP), respectively. In contrast, the third water molecule is not displaced by the NP-A1 isomers and sits only 2.6 Å and 2.2 Å from the methyl groups of NP-A1R and NP-A1S, respectively (**Fig. 3D**).

**Figure 3.**
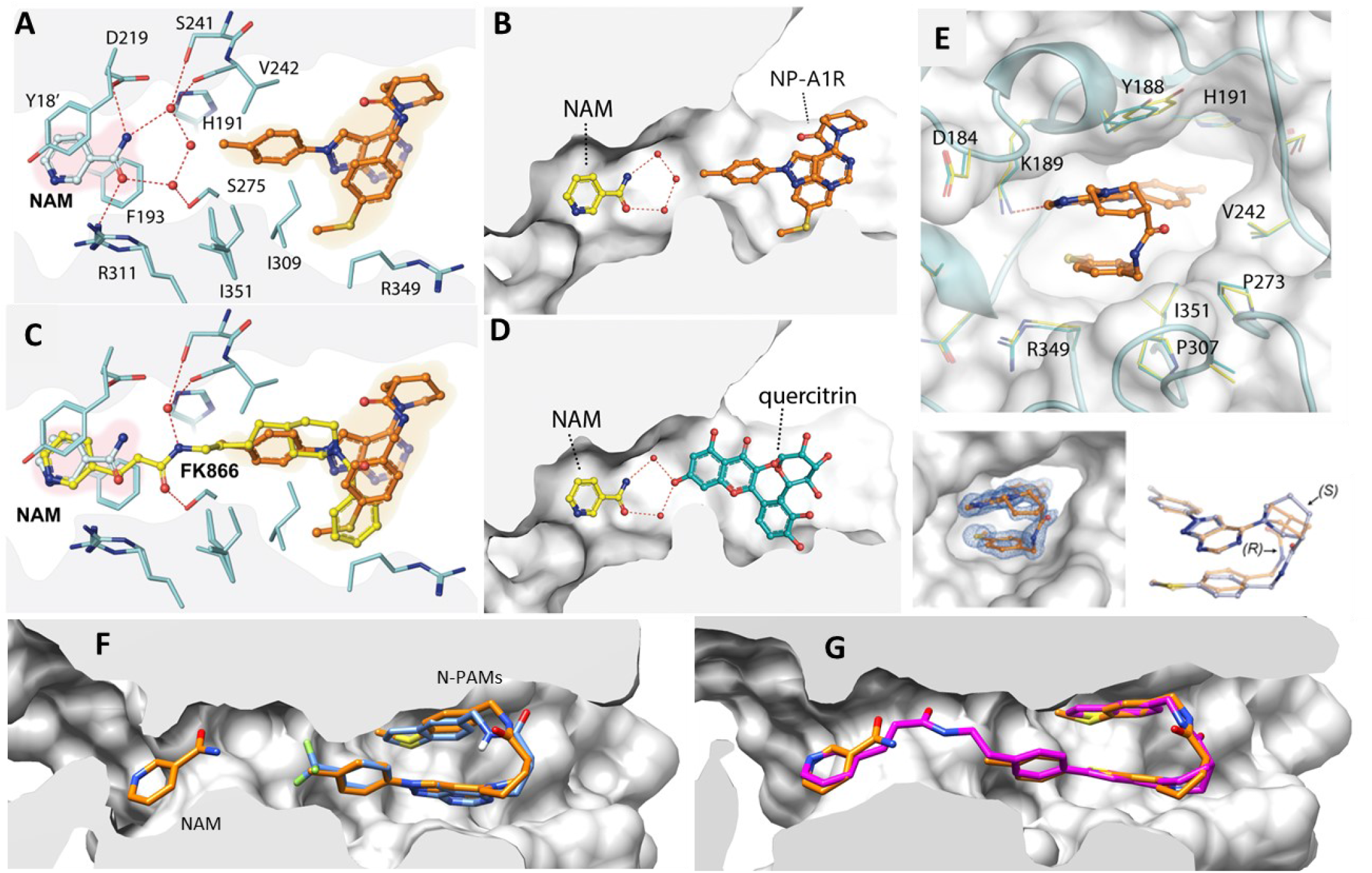
Structures of ligands bound to NAMPT active site and the rear channel. NP-A1R (gold) and NAM showing key residues and H-bonding network (A) and superimposed with bound FK866 (PDB: 2GVJ) (B). Quercitrin (C) or NP-A1R (D) with NAM showing key water molecules. View of bound NP-A1R from mouth of rear channel showing surrounding residues, electron density and superpositio n of S and R isomers extracted from crystal structures (E). NP-A1S (gold) superposed with ZN-2-43 (blue) (F). NPA1S (gold) superpo s ed with NAMPT inhibitor ZN-4-3 (magenta) and NAM.

The replacement of the third water molecule by quercitrin or its apparent compression in the NP-A1 structures migh t be proposed to “push” NAM towards the anomeric carbon of PRPP, potentially compressing the transition state and lowering the activation energy for phosphoribosyl transfer. However, in all co-crystal structures with activator bound, there was no apparent displacement of NAM relative to the position of NAM in structures without activator. Triangulation of the NAM ring-nitrogen, measuring distances to the α-C of residues surrounding the active site (His-247/Arg-392/Asp-354/Asp-313) demonstrated that the position of NAM is identical in the presence or absence of N-PAMs (**Fig. S4**). In crystal structures, neither binding of N-PAM nor quercitrin has any apparent effect on the spatial arrangement of NAM with respect to the α-C of catalytic site amino acid residues, demanding an alternative mechanism of allosteric NAMPT activation.

Crystal structures were examined for other pertinent interactions. Superposition of the structures of quercitrin with NP-A1R shows that the rhamnose ring of quercitrin extends further towards the solvent exposed mouth of the rear channel, with the sugar 2-OH and 3-OH interacting with solvent (**Fig. S5**). The 5-Me group sits in a hydrophobic cleft formed by Pro-307 and Ile-309 and the 4-OH is involved in a H-bond network with the catechol that is itself H-bonded to Lys-189. Comparison with N-PAM cocrystal structures indicates that the H-bonding interaction with Lys-189 is common for these NAMPT activators. Perhaps surprisingly, given its flexibility, the lysine side chain was unperturbed by ligand binding, superimposable in all liganded and unliganded structures (**Fig. S6**).

Both N-PAM isomers bind in the rear channel cavity formed by Tyr-188/Gly-185/Ala-379 and Pro-272/Pro-307, with the tolyl group enclosed by His-191/Val-242/Ile-351 and the benzyl group by Arg-349/Val-350 (**Figs 3E, S6**). In addition to the interaction with Lys-189, N-PAMS form a loose H-bonding network at the mouth of the rear channel: for NP-A1R, this network involves multiple water molecules, the amide and pyrimidine nitrogens of the N-PAM and Thr-304. In addition to facilitating this H-bonding for NP-A1R, the major impact of the piperidine stereocenter is to enforce a twist-boat conformation of the piperidine ring in NP-A1S (**Fig. 3E,F**).

We were not able to obtain a co-crystal structure with the NP-A1 isomers in the absence of bound NAM; however, a structure was obtained with the N-PAM, ZN-2-43, showing that bound NAM is not required for N-PAM binding to the rear channel (**Figs 3F, S7**). ZN-2-43 binds to NAMPT with similar affinity to NP-A1S (**Fig. 2E**). In the absence of a nucleobase, the ZN-2-43 structure has the pi-pi clamp residues in the orientation observed for NAM binding (rather than adenine binding). The same pi-pi clamp orientation is seen in all published co-crystal structures of NAMPT inhibitors and is also observed in the structure of ZN-4-3, a novel NAMPT inhibitor obtained by synthetic extension of the NP-A1 tolyl group with a 5-atom linker to ligate the 3vinylpyridine “warhead” seen in FK866 (**Fig. 1C**). ZN-4 - 3 displays high potency and 5 nM affinity for NAMPT (**Fig. 2D,G**). The co-crystal structure of this N-PAM derived inhibitor bound to the rear channel of NAMPT confirmed the anticipated binding mode with the superposition of the core of ZN-4-3 with the parent N-PAM (**Fig 3G**).

In the three N-PAM co-crystal structures (and the structure of inhibitor ZN-4-3), the ligand adopts an unusual hairpin conformation with intramolecular pi-stacking between the benzyl/benzothiophene and pyrazolo-pyrimidine ring planes (interplane distance ≈ 4.0 Å) (**Figs 3E-G**). In contrast to quercitrin, water molecules are largely excluded from the rear channel, with the hairpin structure filling the rear channel: the volume of the rear channel cavity was calculated as approximately 490-540 Å^3^ (dependent on delineating the mouth of the channel); whereas the volume of the three N-PAMs was calculated to be: 469, 472, and 510 Å^3^; for NP-A1R, NPA1-S, and ZN-2-43, respectively. Intuitively, the hairpin structure suggests an enthalpic cost associated with strain energy; however, DFT molecular orbital calculations show that local minimum energy structures are compatible with the ligand conformations observed in crystal structures (**Fig. S8**). The *R-*isomer accommodates a chair conformation of the piperidine ring in the hairpin structure seen in the rear channel, which translates to high affinity and potency (**Fig. 2;** EC_50_ ≈ 40 nM; K_d_ = 38 nM); whereas the *S-*isomers must adopt a higher energy twist-boat conformation (ΔG = 1.4 kcal/mol) to bind in the rear channel, which translates to lower affinity and potency (**Fig. 2**; EC_50_ ≈ 0.7-1.0 µM; K_d_ = 0.55 µM).

### Mechanism of NAMPT activation

Enzymes have been shown by normal-mode analysis (NMA) (Bahar, Lezon, Bakan, & Shrivastava, 2010) to have open and closed con-formations that correlate with activity; and, ligand binding of an allosteric modulator would be predicted to stabilize conformers that regulate activity.(Khairallah, Ross, & Tastan Bishop, 2021) Using NMA, crystal structures obtained in this work were compared to key NAMPT structures from the literature. The NAMPT structures with BeF_3__coordinated to His-247 provide a mimic of the phospho-H247 phosphoenzyme (PDB: 3DKL; 3DHF; **Fig. 1B**).(Burgos et al., 2009) The structure with an inert analogue of NAM (BzAM), PRPP, and the two Mg^2+^ ions required for catalysis, bound in the active site, can be seen as an ideal model of the enzyme intermediate immediately preceding the transition state for phosphoribosyl transfer to NAM. Comparison of the amplitude (rmsd) of the main - chain residues observed for this structure is indicative of a more “open” protein structure compared to all other structures with bound NAM, which are “closed” (**Fig. S9**). The lowest frequency mode (Mode 1) was only enriched in structures with open conformations. Visualization of this mode demonstrates coupling of N-PAM binding in the rear channel to residues remote from the rear channel, indicating a potential allosteric mechanism mechanically coupling rear channel binding to the active site (**Fig. S9**).

NAMPT uses cellular ATP to drive the salvage of NAM to synthesize NMN. The catalytic efficiency (k_cat_/K_M_) of NAMPT results from the ATPase reaction, generating phospho-H247. This lowers the K_M_ for PRPP and consequently lowers K_M_(NAM) 170 fold, while increasing k_cat_ almost 20-fold.(Burgos & Schramm, 2008) In the absence of ATP, the equilibrium established by NAMPT (PRPP + NAM ⇋NMN + PPi) lies to the left, as demonstrated clearly by the conversion of NMN to NAM seen in the absence of ATP (**Fig. 4A**). Although NAMPT does not experience zero ATP in cells, this conceptual experiment is useful to emphasize the dependence on ATP. Addition of ATP shifts the equilibrium towards NMN, demonstrating the importance of ATP-mediated formation of phospho-H247 for the forward reaction. In the presence of NP-A1S or SBI-0797812, the equilibrium remains in favor of NAM in the absence of ATP and shifts towards NMN on addition of ATP (2 mM) (**Fig. 4A**). Similar experiments either without NAM, or without NMN in the incubation mixture, reinforce the importance of ATP: N-PAM activation of NAMPT is ATP-dependent (**Fig. S10**).

**Figure 4.**
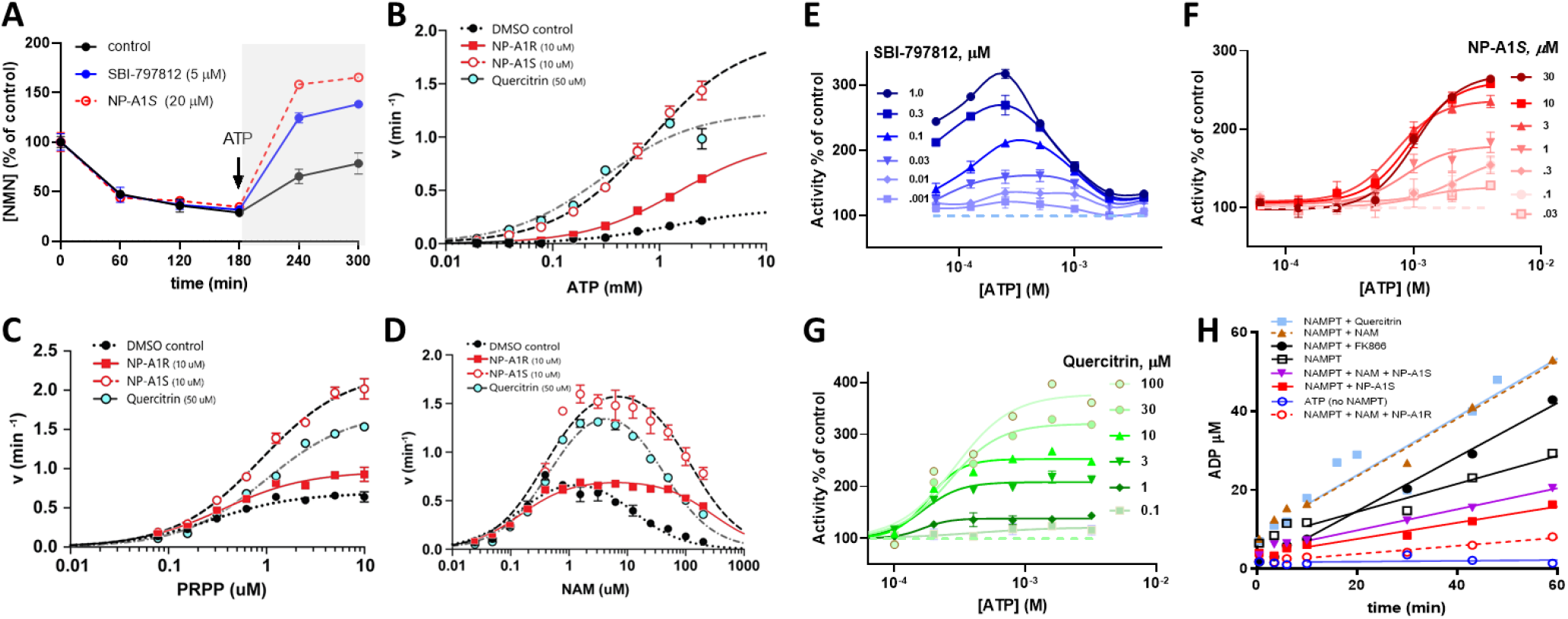
NAMPT dependence on ATP and NAM. Dependence on ATP of equilibrium shift from NAM to NMN in presence or abs en ce of activators (A). Dependence of activity on ATP (B), PRPP (C), and NAM (D) for N-PAMs and quercitrin. Dependence of activ ity o n ATP for SBI-797812 (E), NP-A1S (F), and quercitrin (G. Regulation of uncoupled ATPase reaction is ligand dependent (H).

### Activity dependence on NAM, PRPP, and ATP

In the primary enzyme assay N-PAMs and quercitrin increased rate without significant effect on K_M_(ATP); and a similar observation was made for K_M_(PRPP) (**Figs 4B,C; Table S1**). NP-A1S increased catalytic efficiency (V_max_/K_M_) with respect to ATP tenfold (**Table S1**). Under the conditions studied, the K_M_ for NAM (≈145 nM) was not influenced by NP-A1R and modestly increased for NP-A1S; whereas the K_I_ of NAM was significantly right-shifted by N-PAMs by 10-to 20-fold (**Fig. 4D; Table S1**). The attenuation by N-PA M of substrate inhibition by NAM is significant, since this opens a window of N-PAM-stimulated NAMPT activity extending to 500 µM NAM, which does not exist for the enzyme in the absence of N-PAM.

The orthogonal enzyme assay was used to compare the ATP dependence of SBI-0797812 activation with that of NP-A1S and quercitrin. Although K_M_(ATP) is reported as 7 mM, substrate inhibition by [ATP] > 4 mM, is also reported (Burgos & Schramm, 2008). The 4-pyridyl activator, SBI-0797812, caused a significant left-shift in K_M_(ATP) and also drastically left-shifted the K_I_(ATP) to < 1mM (**Fig. 4E**). As seen in the coupled enzyme assay (**Fig. 4B**), NP-A1S had no significant effect on ATP dependence (**Fig. 4F**) and a similar observation was made for quercitrin (**Fig. 5C**) and a related phenolic activator (**Fig. S11**). The ATP dependence observed for SBI-0797812 suggests activation will be highly dependent on ATP concentration in a cellular context. Physiological, cellular ATP concentrations vary from 2-7 × 10^−3^ M, depending on cell type(Greiner & Glonek, 2021); whereas the concentration of ATP in plasma is 10,000-fold lower.(Gorman, Feigl, & Buffington, 2007) The optimal activation driven by SBI-0797812 is therefore predicted to lie between ATP concentrations experienced by cellular NAMPT and extracellular NAMPT (eNAMPT).

**Figure 5.**
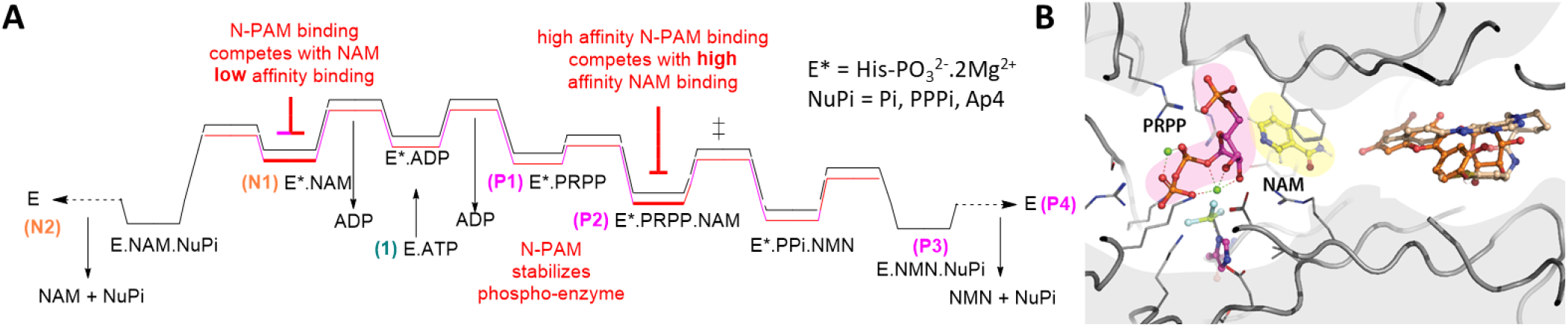
N-PAM mechanism of action. A) Stepwise mechanism starting with (1) ATP binding and ATPase reaction to give phospho- enzyme: Non-productive pathway; (N1) Low affinity binding of NAM; leading to (N2) uncoupled, non - productive ATPase reaction; Productive pathway: (P1) PRPP binding leads to; (P2) high affinity NAM binding and subsequent turnover to NMN, followed by; (P3) breakdown of the phospho-enzyme by capture of metaphosphate (PO_3__) by water, PPi, or ATP (Amici et al., 2017; Thatch er & Klu ger, 1989), leaving NAMPT poised for “reloading” by ATP. Red text describes modulation by N-PAM. B) Structure of NAMPT activ e s ite containing PRPP and phospho-H247, nucleobase pocket containing NAM, and rear channel containing N-PAM or quercitrin. In the p ro posed mechanism, NAM binding occurs via the rear channel.

The ATPase and NAM salvage reactions catalyzed by NAMPT are uncoupled (or non-stoichiometric), resulting in futile consumption of ATP: i.e. the ratio of ADP/NMN formation is greater than unity under most conditions. (Burgos & Schramm, 2008) For example, incubating with ATP (2 mM) for 60 min, Burgos et al. reported formation of 30 µM ADP, which was reduced in the presence of NAM+PRPP or NMN+PPi, and increased fourfold in the presence of NAM alone and sevenfold in the presence of PPi alone.(Burgos & Schramm, 2008) Thus, the binding of NAM in the absence of PRPP leads to accelerated non-productive ATP degradation to ADP. Published results with NAM were recapitulated and quercitrin was also observed to increase uncoupled ATP consumption (**Fig. 4G**). Addition of the NAMPT inhibitor, FK666, also increased ATP turnover to the same high rate observed for NAM (for the inhibitor, the burst of ADP, seen for NAM was, not observed). This is consistent with accelerated breakdown of the phospho-enzyme when a ligand is bound to the nucle-obase pocket in the absence of PRPP. The N-PAMs reduced ATP consumption in the presence of NAM, with the high affinity NP-A1R more effective than the lower affinity *S-*isomer. The N-PAMs also limited ATP consumption in the absence of NAM. Thus N-PAMs inhibit non-productive binding of NAM and are diametrically differentiated from the phenolic activators in regulation of the ATPase reaction catalyzed by NAMPT.

### The importance of the rear channel in NAMPT function

The detailed enzyme mechanism elegantly delineated by Schramm, Burgos and co-workers does not reference the rear channel.(Burgos et al., 2009; Burgos & Schramm, 2008; Burgos et al., 2013) An extended mechanism to account for N-PAM activation must take account of the previously detailed mechanism and the role of the rear channel (**Fig. 5**). Applying Occam’s razor, we propose a mechanism that, like Schramm’s, relies primarily on modulation of NAM binding with an additional contribution to k_cat_. The proposed reaction profile (**Fig. 5A**) incorporates nonproductive low affinity (high K_M_) NAM binding leading to ATP catabolism and high affinity (low K_M_) NAM binding leading to turnover to NMN. High affinity NAM binding occurs, reasonably, via the rear channel, since PRPP is required to be bound to the phosphoenzyme and we propose that low affinity NAM binding is also via the rear channel.

In this scenario, N-PAM binding inhibits low affinity NAM binding, shifting K_I_(NAM) to the right, essentially derepressing the effect of higher [NAM] on enzyme activity. In the case of the high affinity N-PAM *R-*isomer, high affinity NAM binding is also partially inhibited, leading to lower fold-activation.

We propose a smaller contribution from N-PAM stabilization of the “open” states, containing the phosphoenzyme (E*), as suggested by NMA. The phosphoenzyme sits adjacent to the transition states for ribosylation and dephosphorylation; therefore, allosteric stabilization by N - PAM binding will lower the activation barriers for these reactions, translating to an increase in k_cat_. The proposed mechanism is driven by modulation of NAM binding and supported by: i) the attenuation of NAM-induced uncoupling of the ATPase reaction (**Fig. 4H**); ii) the differences in potency and activation of turnover by the two NP-A1 isomers (**Figs 2C,E**); and iii) the observed >20-fold right-shift in K_I_(NAM) observed for N-PAMs (**Fig. 4D**).

Access of other reaction components via the rear channel is speculative; however, it is noted that phosphoribosylated inhibitors, such as GNE-617-PR (**Fig. 1C**) bind to NAMPT reversibly.(Oh et al., 2014)

### Regulation of cellular NAD^+^

Given the dependence of NAMPT activation on NAM, ATP, and NAD^+^ concentrations, the translation of the actions of a biochemical activator to a cellular context is not guaranteed. Using FK-866 and NMN as negative and positive controls, respectively, we explored several cell lines to select a highly reproducible model system with high dynamic range. We optimized a commercial NADglo assay in the THP-1human leukemic monocyte cell line, measuring NAD^+^ after incubation with test compounds for 24 h. Cells treated with NP-A1S responded with an increase in cellular NAD^+^ of over twofold, whereas NP-A1R only increased cellular NAD^+^ 1.2-fold (**Fig. 6A**). The N-PAM, ZN-2-43S, was more potent and more efficacious than NP-A1R. Quercitrin was inactive and a recently reported example of a phenolic NAMPT activator, NAT1 (**Fig. 1C**), increased cellular NAD^+^ 1.25-fold (**Fig. 7A**). Surprisingly, the recently reported NAMPT activator, SBI-0797812, reduced cellular levels of NAD^+^ in THP-1 cells (**Fig. 6A**). To explore a rationale for this observation, we re-examined the biochemical activation of NAMPT at a higher, physiologically relevant ATP concentration (4 mM) and high and low NAM concentrations (3 µM is at the lower end of plasma NAM levels(Liu et al., 2018)). Inhibition of NAMPT activity was observed for SBI-079812 (**Fig. 7**), compatible with observations on ATP dependence and cellular NAD^+^.

**Figure 6.**
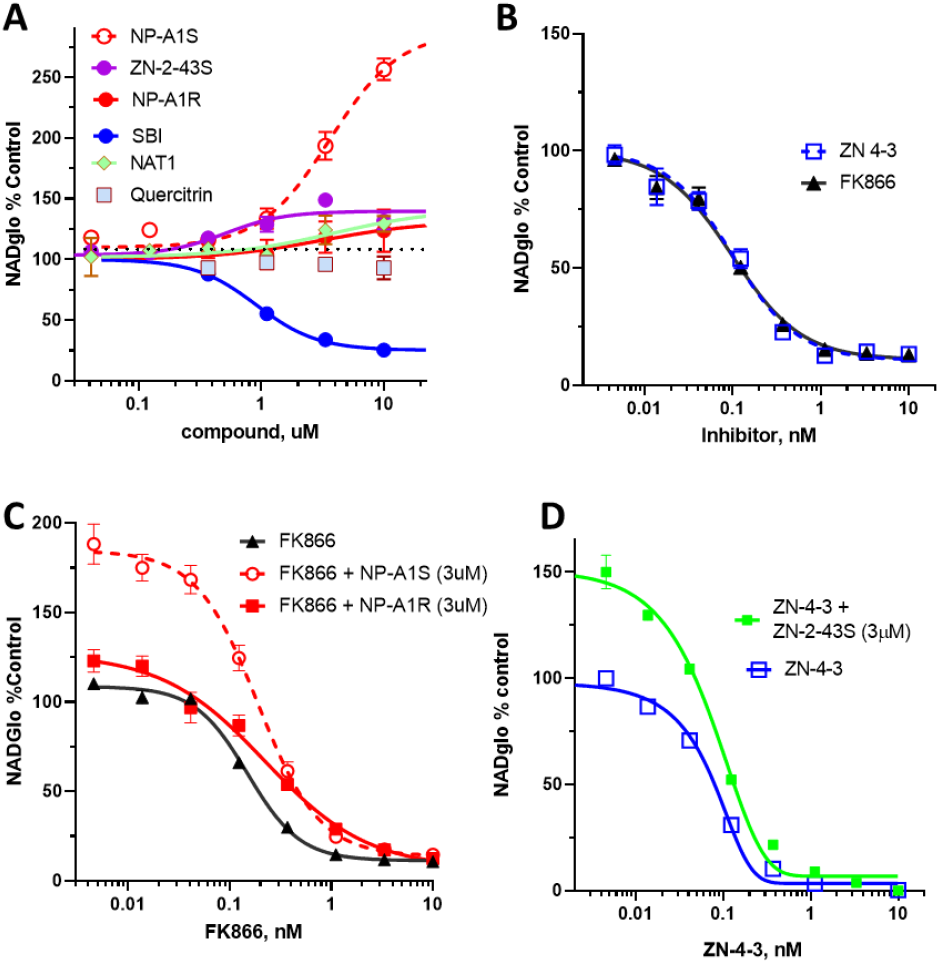
NAD^+^ regulation in THP-1 cells. Response to: (A) N- PAMs and activators and (B) NAMPT inhibitors. The concentration response for inhibition by FK866 was right-shifted by N-PAM (C) and for ZN-4-3 was right-shifted by N-PAM (D).

**Figure 7.**
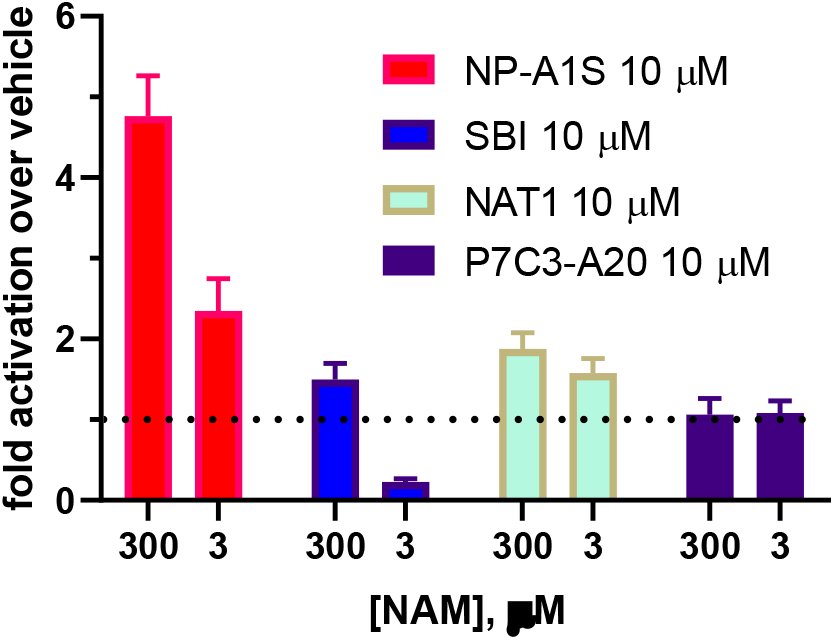
NAMPT activation by N-PAMs compared to other reported activators. Activity relative to vehicle control was measured in the orthogonal enzyme assay: ATP 4 mM; NAM 3 or 30 µM; PRPP 40 µM.

For N-PAMs, the biochemical observations on NAMPT activation fully translated to modulation of NAD^+^ in a cellular context (**Fig. 6A**). In addition to NAMPT inhibition by A TP and NAM, the reported K_I_ for inhibition of NAMPT activity by NAD^+^ is 2 µM;(Burgos & Schramm, 2008) therefore, NAMPT activity was measured in the presence of N-PAMs at 10 µM NAD^+^, showing retention of NAMPT activity at a level equal or superior to that in the absence of NAD^+^ (**Fig. S12**).

It was straightforward synthetically to convert an N-PAM to a novel chemical class of NAMPT inhibitor: the potent biochemical inhibition of NAMPT by ZN-4-3 translated to picomolar potency in cells, *in simile* with FK866 (IC_50_ = 100 pM; **Fig. 6B**). Having demonstrated that N-PAMs bind to the same channel as NAMPT inhibitors such as ZN-4-3 and FK-866, we tested the ability of N-PAMs to compete with and displace inhibitors in THP-1 cells. Treatment of cells with both NP-A1 isomers in the presence of FK-866 produced a right-shift in the FK-866 response (to IC_50_ = 0.19 nM and 0.22 nM for *S* and *R* isomers, respectively) (**Fig. 6C**). The effect of the high affinity *R-*isomer is compatible with its ability to compete with FK866 for binding to NAMPT, but with inherently modest activation of NAMPT.

Biochemical activity does not necessarily translate directly to activity in cells, even with good cell permeability, as has been observed for NAMPT inhibitors.(Oh et al., 2014) The cellular actions of N-PAM, ZN-2-43, and NAMPT inhibitor, ZN-4-3, were as predicted from biochemical observations (**Fig. 6D**). N-PAMs increased cellular NAD^+^ levels: the *S-*isomer increasing cellular NAD^+^ by 250%; and the *R-*isomer blocking of the inhibitory activity of FK866, even though NAMPT activation was modest.

Although quercitrin was inactive in cells, the phenol, NAT1, acted as a NAMPT activator in cells and biochemical assays (**Figs. 6A, 7**). The observation that several biogenic and/or bioactive phenols are NAMPT activators may indicate that this activity contributes to their observed biological actions; although, it is unsafe to ascribe the in vivo phenotype of such phenols entirely to NAMPT activation. The reported in vivo phenotype of NAT1 includes neuroprotection;(Yao et al., 2022) however, such phenols have other validated targets and are classical antioxidants providing multiple mechanisms of neuroprotection.(Abdelhamid et al., 2011) In multiple reports, P7C3-A20 continues to be studied as a NAMPT activator, its pharmacological effects, including neuroprotection,(Pieper & McKnight, 2019) consistent with those of an agent that elevates cellular NAD^+^ (Fig. **S13**).(Gardell et al., 2019) P7C3-A20 does not increase NAMPT activity in biochemical assays (**Fig. 7**).

## CONCLUSIONS

NAMPT is proposed to play multiple physiological roles in addition to being the rate-limiting enzyme in mammalian NAD^+^ biosynthesis. NAMPT is proposed to act as a cytokine sometimes known as pre-B cell colony-enhancing factor (PBEF),(Rongvaux et al., 2002; Samal et al., 1994) and as an adipokine known as visfatin.(Hug & Lodish, 2005) In its guise as a cytokine, extracellular NAMPT (e NAMPT) acts as a ligand and agonist at the Toll-Like Receptor 4 (TLR4) activating inflammatory responses.(Camp et al., 2015) Much remains to be discovered about the interplay between eNAMPT and cellular NAMPT, the latter controlling salvage of NAM. These roles are pivotal to cellular bioenergetics, mitochondrial function, metabolic homeostasis, and the Circadian clock.

Small molecules were identified that increase the catalytic efficiency of NAMPT for NAM salvage. Novel NAMPT positive allosteric modulators (N-PAMs) were differentiated from phenolic activators, represented by quercitrin and NAT1, and recently reported 4-pyridyl activators, represented by SBI-0797812 (**Fig. 7**). Differentiation was observed in dependence on ATP and NAM concentration an d modulation of the uncoupled ATPase reaction of NAMPT. The observations on quercitrin and other biogenic and/ or bioactive phenols suggest that, at higher micromolar concentrations, NAMPT activation may contribute to the biological phenotype of these and related compounds and natural products.

Cellular NAD^+^ biosynthesis is tightly regulated; notably via feedback inhibition by ATP, NAM, and NAD^+^. A key aspect of the differentiation of activators is associated with this inhibition by ATP, NAM, and NAD^+^. SBI-0797812 induced a dramatic left-shift in both K_M_ and K_I_ for ATP and this activator *inhibited* both NAMPT activity and NAD^+^ formation in THP-1 cells. Although the K_M_ for ATP and NAM was not perturbed by N-PAMs, the right-shift in K_I_(NAM) results in enhanced enzyme activity at higher cellular concentrations of NAM. Elevated NAM levels (> K_I_), resulting from accelerated NAD^+^ catabolism in stressed cells would normally induce a paradoxical feedback inhibition of NAMPT; however, N-PAM binding to the rear channel of NAMPT was shown to relieve this inhibition, promoting NAD^+^ biosynthesis. Importantly, biochemical observations on NAMPT activity translated to regulation of cellular NAD^+^, measured in THP-1 cells. Taken together, these observations indicate that NAMPT activators can be designed with tissue and cell selectivity, dependent on cellular concentrations of substrates and products.

The mechanism of N-PAM activation was determined by synthesis of novel activators, inhibitors, and an FP-probe. Using multiple co-crystal structures and enzyme kinetics, a mechanism is proposed in which N-PAM binding to the rear channel competes with NAM access to the nucleobase pocket via this channel. Maximal activation is induced by an N-PAM with affinity for the rear channel that allows productive (low K_M_) NAM binding and blocks nonproductive (high K_M_) binding (**Fig. 5**). To account for a modest increase in k_cat_, N-PAMs are proposed to stabilize the phospho-enzyme through allosteric binding remote (≈ 20Å) from phospho-His-247 through mechanical coupling to the active site. This composite mechanism is compatible with the observed coupling of ATP utilization to NAM turnover by N-PAMs. Central to the mechanism of activation by N-PAMs and phenolic activators is the rear channel, a unique feature of NAMPT, raising the question as to whether endogenous regulators exist that bind to this structural feature.

Pharmacological activation of NAMPT is a promising therapeutic strategy, in metabolic disorders and diseases of aging, to elevate levels of cellular NMN and NAD^+^ (Parihar & Brewer, 2007; Revollo et al., 2007; Rongvaux et al., 2002; J. Yoshino et al., 2011; Zhu et al., 2015). The research presented herein provides a mechanism for pharmacological modulation of NAMPT, which given the dependence on cellular concentrations of ATP, NAM, and NAD^+^, is predictive of cell and tissue selectivity that itself can be exploited therapeutically.

## Materials and Methods

For full materials and methods, please see *Supplemental Information*.

## Chemicals, Reagents, and Assays

Synthesis and characterization for all compounds, experimental details of biochemical, biophysical, and cellular assays, and supplemental table and figures. The X-ray coordinates have been deposited with the Protein Data Bank.

## Grant Support

This study is supported by NIH grant RF1AG067771. JG was supported, in part, by NIH T32AG57468 and MSL by NIH T32 GM008804. This research used resources of the Advanced Photon Source, a U.S. Department of Energy (DOE) Office of Science User Facility operated for the DOE Office of Science by Argonne National Laboratory under Contract No. DE - AC 02 - 06CH11357. Use of the LS-CAT Sector 21 was supported by the Michigan Economic Development Corporation and the Michigan Technology Tri-Corridor (Grant 085P1000817).

### Abbreviations

ADH: alcohol dehydrogenase
ADRD: Alzheimer’s disease and related dementia
ARTs: ADP-ribosyltransferases
eNAMPT: extracellular NAMPT
FP: fluorescence polarization
MST: microscale thermophoresis
N-PAMs: NAMPT positive allosteric modulators
NAD^+^: nicotinamide adenine dinucleotide
NADP^+^: 2-phosphate derivative
NAM: nicotinamide
NAMPT: nicoti n amide phosphoribosyl-transferase
NAPRT: nicotinic acid phosphoribosyltransferase
NMA: normal - mode analysis
NMN: nico - tinamide mononucleotide
NMNAT: nicotinamide/nicotinic acid mononucleotide adenylyltransferase
PARP: poly - ADP- r i bo se polymerase
phosphor-H247: *N-*phosphohistidine
PDB: protein data bank
PRPP: α-D-5-phosphoribosyl-1- py ro phos ph ate
SARM1: sterile alpha and TIR motif containing 1
SIRTs: sirtuins
TLR4: Toll-Like Receptor 4
T2D: type-2 diabetes.

## Notes

**Disclosures:** The authors have no conflicts to disclose. G.T. is an inventor on patents owned by the University of Illinois.

### Competing Interest Statement

G.T. is an inventor on patents owned by the University of Illinois.
This statement is required by a COI management plan

## REFERENCES

Abdelhamid, R., Luo, J., Vandevrede, L., Kundu, I., Michalsen, B., Litosh, V. A., … Thatcher, G. R. (2011). Benzothiophen e Selective Estrogen Receptor Modulators Provide Neuroprotection by a novel GPR30-dependent Mechanism. ACS Chem Neurosci, 2(5), 256–268. doi:10.1021/cn100106a

Akiu, M., Tsuji, T., Iida, K., Sogawa, Y., Terayama, K., Yokoyama, M., … Nakamura, T. (2021). Discovery of DS68702229 as a Potent, Orally Available NAMPT (Nicotinamide Phosphoribosyltransferase) Activator. Chem Pharm Bull (Tokyo), 69(11), 1110–1122. doi:10.1248/cpb.c21-00700

Amici, A., Grolla, A. A., Del Grosso, E., Bellini, R., Bianchi, M., Travelli, C., … Orso mando, G. (2017). Synthesis and degradation of adenosine 5’-tetraphosphate by nicotinamide and nicotinate phosphoribosyltransferases. Cell Chem Biol, 24(5), 553–564 e554. doi:10.1016/j.chembiol.2017.03.010

Bahar, I., Lezon, T. R., Bakan, A., & Shrivastava, I. H. (2010). Normal mode analysis of biomolecular structures: functional mechanisms of membrane proteins. Chem Rev, 110(3), 1463–1497. doi:10.1021/cr900095e

Burgos, E. S., Ho, M. C., Almo, S. C., & Schramm, V. L. (2009). A phosphoenzyme mimic, overlappin g catalytic sites and reaction coordinate motion for human NAMPT. Proc Natl Acad Sci U S A, 106(33), 13748–13753. doi:10.1073/pnas.0903898106

Burgos, E. S., & Schramm, V. L. (2008). Weak coupling of ATP hydrolysis to the chemical equilibrium of human nicot inami de phosphoribosyltransferase. Biochemistry, 47(42), 11086–11096. doi:10.1021/bi801198m

Burgos, E. S., Vetticatt, M. J., & Schramm, V. L. (2013). Recycling nicotinamide. The transition -state structure of human nicotinamide phosphoribosyltransferase. J Am Chem Soc, 135(9), 3485–3493. doi:10.1021/ja310180c

Camacho-Pereira, J., Tarrago, M. G., Chini, C. C. S., Nin, V., Escande, C., Warner, G. M., … Chini, E. N. (2016). CD38 dictates age-related NAD decline and mitochondrial dysfunction through an SIRT3-dependent mechanism. Cell Metab, 23(6), 1127–1139. doi:10.1016/j.cmet.2016.05.006

Camp, S. M., Ceco, E., Evenoski, C. L., Danilov, S. M., Zhou, T., Chiang, E. T., … Garcia, J. G. (2015). Unique Toll –Like Receptor 4 Activation by NAMPT/PBEF Induces NFkappaB Signaling and Inflammatory Lung Injury. Sci Rep, 5, 13135. doi:10.1038/srep13135

Chini, C. C. S., Peclat, T. R., Warner, G. M., Kashyap, S., Espindola-Netto, J. M., de Oliveira, G. C., … Chini, E. N. (2020). CD38 ecto-enzyme in immune cells is induced during aging and regulates NAD(+) and NMN levels. Nat Metab, 2(11), 1284–1304. doi:10.1038/s42255-020-00298-z

Chowdhry, S., Zanca, C., Rajkumar, U., Koga, T., Diao, Y., Raviram, R., … Mischel, P. S. (2019). NAD metabolic dependency in cancer is shaped by gene amplification and enhancer remodelling. Nature, 569(7757), 570–575. doi:10.1038/s41586-019-1150-2

Dellinger, R. W., Santos, S. R., Morris, M., Evans, M., Alminana, D., Guarente, L., & Marcotulli, E. (2017). Repeat dose NRPT (nicotinamide riboside and pterostilbene) increases NAD(+) levels in humans safely and sustainably: a rand o mi zed, double-blind, placebo-controlled study. NPJ Aging Mech Dis, 3, 17. doi:10.1038/s41514-017-0016-9

Essuman, K., Summers, D. W., Sasaki, Y., Mao, X., DiAntonio, A., & Milbrandt, J. (2017). The SARM1 Toll/Interleukin -1 receptor domain possesses intrinsic NAD(+) cleavage activity that promotes pathological axonal degeneration. Neuron, 93(6), 1334–1343 e1335. doi:10.1016/j.neuron.2017.02.022

Fang, E. F., Scheibye-Knudsen, M., Brace, L. E., Kassahun, H., SenGupta, T., Nilsen, H., … Bohr, V. A. (2014). Defective mitophagy in XPA via PARP-1 hyperactivation and NAD(+)/SIRT1 reduction. Cell, 157(4), 882–896. doi:10.1016/j.cell.2014.03.026

Gardell, S. J., Hopf, M., Khan, A., Dispagna, M., Hampton Sessions, E., Falter, R., … Pinkerton, A. B. (2019). Boosting NAD(+) with a small molecule that activates NAMPT. Nat Commun, 10(1), 3241. doi:10.1038/s41467-019-11078-z

Garten, A., Schuster, S., Penke, M., Gorski, T., de Giorgis, T., & Kiess, W. (2015). Physiological and pathophysiological rol es of NAMPT and NAD metabolism. Nat Rev Endocrinol, 11(9), 535–546. doi:10.1038/nrendo.2015.117

Gomes, A. P., Price, N. L., Ling, A. J., Moslehi, J. J., Montgomery, M. K., Rajman, L., … Sinclair, D. A. (2013). Declining NAD(+) induces a pseudohypoxic state disrupting nuclear-mitochondrial communication during aging. Cell, 155 (7), 1624–1638. doi:10.1016/j.cell.2013.11.037

Gordon-Blake, J. M., Karumudi, B., Ratia, K., Knopp, R. C., Dye, K., Ben Aissa, M., & Thatcher, G. R. J. (2019). Novel NAMPT activators attenaute neurotoxicity and neuroinflammation associated with neurodegeneration. Alzheimer’s Dementia, 15(75), 266–267. doi:10.1016/j.jalz.2019.06.108

Gorman, M. W., Feigl, E. O., & Buffington, C. W. (2007). Human plasma ATP concentration. Clin Chem, 53(2), 318–325. doi:10.1373/clinchem.2006.076364

Greiner, J. V., & Glonek, T. (2021). Intracellular ATP Concentration and Implication for Cellular Evolution. Biology (Basel), 10(11). doi:10.3390/biology10111166

Guarente, L. (2014). Linking DNA damage, NAD(+)/SIRT1, and aging. Cell Metab, 20(5), 706–707. doi:10.1016/j.cmet.2014.10.015

Hug, C., & Lodish, H. F. (2005). Medicine. Visfatin: a new adipokine. Science, 307(5708), 366–367. doi:10.1126/science.1106933

Imai, S., & Guarente, L. (2014). NAD+ and sirtuins in aging and disease. Trends Cell Biol, 24(8), 464–471. doi:10.1016/j.tcb.2014.04.002

Khairallah, A., Ross, C. J., & Tastan Bishop, O. (2021). GTP Cyclohydrolase I as a Potential Drug Target: New Insights into Its Allosteric Modulation via Normal Mode Analysis. J Chem Inf Model, 61(9), 4701–4719. doi:10.1021/acs.jcim.1c00898

Liu, L., Su, X., Quinn, W. J., 3rd, Hui, S., Krukenberg, K., Frederick, D. W., … Rabinowitz, J. D. (2018). Quantitative Analysis of NAD Synthesis-Breakdown Fluxes. Cell Metab, 27(5), 1067–1080 e1065. doi:10.1016/j.cmet.2018.03.018

Martens, C. R., Denman, B. A., Mazzo, M. R., Armstrong, M. L., Reisdorph, N., McQueen, M. B., … Seals, D. R. (2018). Chronic nicotinamide riboside supplementation is well-tolerated and elevates NAD(+) in health y mi d d l e - aged an d older adults. Nat Commun, 9(1), 1286. doi:10.1038/s41467-018-03421-7

Mitchell, S. J., Bernier, M., Aon, M. A., Cortassa, S., Kim, E. Y., Fang, E. F., … de Cabo, R. (2018). Nicotinamide impro ves aspects of healthspan, but not lifespan, in mice. Cell Metab, 27(3), 667–676 e664. doi:10.1016/j.cmet.2018.02.001

Oh, A., Ho, Y. C., Zak, M., Liu, Y., Chen, X., Yuen, P. W., … Wang, W. (2014). Structural and biochemical analyses of the catalysis and potency impact of inhibitor phosphoribosylation by human nicotinamide phosphoribo s yl trans feras e. Chembiochem, 15(8), 1121–1130. doi:10.1002/cbic.201402023

Parihar, M. S., & Brewer, G. J. (2007). Mitoenergetic failure in Alzheimer disease. Am J Physiol Cell Physiol, 292(1), C8–23. doi:10.1152/ajpcell.00232.2006

Pieper, A. A., & McKnight, S. L. (2019). Benefits of Enhancing Nicotinamide Adenine Dinucleotide Levels in Damaged or Diseased Nerve Cells. Cold Spring Harbor Symposia on Quantitative Biology. doi:10.1101/sqb.2018.83.037622

Revollo, J. R., Korner, A., Mills, K. F., Satoh, A., Wang, T., Garten, A., … Imai, S. (2007). Nampt/PBEF/Visfatin regulates insulin secretion in beta cells as a systemic NAD biosynthetic enzyme. Cell Metab, 6(5), 363–375. doi:10.1016/j.cmet.2007.09.003

Rongvaux, A., Shea, R. J., Mulks, M. H., Gigot, D., Urbain, J., Leo, O., & Andris, F. (2002). Pre-B-cell colony-enhancing factor, whose expression is up-regulated in activated lymphocytes, is a nicotinamide phosphoribosy l tran s feras e, a cytosolic enzyme involved in NAD biosynthesis. Eur J Immunol, 32(11), 3225–3234. doi:10.1002/1521-4141(200211)32:11<3225::AID-IMMU3225>3.0.CO;2-L

Samal, B., Sun, Y., Stearns, G., Xie, C., Suggs, S., & McNiece, I. (1994). Cloning and characterization of the cDNA encoding a novel human pre-B-cell colony-enhancing factor. Mol Cell Biol, 14(2), 1431–1437.

Stromsdorfer, K. L., Yamaguchi, S., Yoon, M. J., Moseley, A. C., Franczyk, M. P., Kelly, S. C., … Yoshino, J. (2016). NAMPT-Mediated NAD(+) Biosynthesis in Adipocytes Regulates Adipose Tissue Function and Multi -organ Insulin Sensitivity in Mice. Cell Rep, 16(7), 1851–1860. doi:10.1016/j.celrep.2016.07.027

Thatcher, G. R. J., & Kluger, R. (1989). Mechanism and catalysis of nucleophilic substitution in phosphate esters. In D. Beth ell (Ed.), Adv Phys Org Chem (Vol. 25, pp. 99–265).

Uddin, G. M., Youngson, N. A., Doyle, B. M., Sinclair, D. A., & Morris, M. J. (2017). Nicotinamide mononucleotide (NMN) supplementation ameliorates the impact of maternal obesity in mice: comparison with exercise. Sci Rep, 7(1), 15063. doi:10.1038/s41598-017-14866-z

Wilsbacher, J. L., Cheng, M., Cheng, D., Trammell, S. A. J., Shi, Y., Guo, J., … Tse, C. (2017). Discovery and Characterization of Novel Nonsubstrate and Substrate NAMPT Inhibitors. Mol Cancer Ther, 16(7), 1236–1245. doi:10.1158/1535-7163.MCT-16-0819

Yao, H., Liu, M., Wang, L., Zu, Y., Wu, C., Li, C., … Wang, G. (2022). Discovery of small -molecule activators of nicotinami de phosphoribosyltransferase (NAMPT) and their preclinical neuroprotective activity. Cell Res, 32(6), 570–584. doi:10.1038/s41422-022-00651-9

Yoshino, J., Baur, J. A., & Imai, S. I. (2018). NAD(+) intermediates: the biology and therapeutic potential of NMN and NR. Cell Metab, 27(3), 513–528. doi:10.1016/j.cmet.2017.11.002

Yoshino, J., Mills, K. F., Yoon, M. J., & Imai, S. (2011). Nicotinamide mononucleotide, a key NAD(+) intermediate, treats the pathophysiology of diet- and age-induced diabetes in mice. Cell Metab, 14(4), 528–536. doi:10.1016/j.cmet.2011.08.014

Yoshino, M., Yoshino, J., Kayser, B. D., Patti, G. J., Franczyk, M. P., Mills, K. F., … Klein, S. (2021). Nicotinamide mononucleotide increases muscle insulin sensitivity in prediabetic women. Science, 372(6547), 1224–1229. doi:10.1126/science.abe9985

Zhang, R. Y., Qin, Y., Lv, X. Q., Wang, P., Xu, T. Y., Zhang, L., & Miao, C. Y. (2011). A fluorometric assay for high -throughput screening targeting nicotinamide phosphoribosyltransferase. Anal Biochem, 412(1), 18–25. doi:10.1016/j.ab.2010.12.035

Zheng, X., Bauer, P., Baumeister, T., Buckmelter, A. J., Caligiuri, M., Clodfelter, K. H., … Bair, K. W. (2013). Structure-based identification of ureas as novel nicotinamide phosphoribosyltransferase (Nampt) inhibitors. J Med Chem, 56(12), 4921–4937. doi:10.1021/jm400186h

Zhu, X. H., Lu, M., Lee, B. Y., Ugurbil, K., & Chen, W. (2015). In vivo NAD assay reveals the intracellular NAD contents and redox state in healthy human brain and their age dependences. Proc Natl Acad Sci U S A, 112(9), 2876–2881. doi:10.1073/pnas.1417921112

