## Supplemental Information for "The mechanism of nicotinamide phosphoribosyltransferase whereby positive allosteric modulation elevates cellular NAD^+^"

### MATERIALS & METHODS

#### BIOCHEMISTRY

*Purification of NAMPT enzyme*: pET21a vector with human wild-type NAMPT in frame with a C-terminal $His_6$-tag was obtained from Genscript. NAMPT protein expression was induced using an auto-induction medium described.(1) Cells were grown for 24h at 20°C, harvested by centrifugation, and lysed by sonication in Buffer A (20 mM Tris-HCl, pH 8, 0.5 M NaCl, 8 mM imidazole) containing Roche EDTA-free protease inhibitor tablets, 10 ug/mL DNaseI, and 10 ug/mL lysozyme. The lysate was clarified by centrifugation and loaded onto a 5-mL Talon cobalt column equilibrated with Buffer A. NAMPT protein was eluted with Buffer B (20 mM Tris-HCl, pH 8.0, 0.5 M NaCl, 50 mM imidazole). Fractions containing protein were pooled, concentrated, and exchanged into 20 mM Tris-HCl, pH 8, 150 mM NaCl, 5 mM DTT and used directly for crystallization or flash frozen and stored at -80°C.

*NAMPT coupled enzyme activity assay***.** The NAMPT primary enzyme assay is based on conditions from Burgos and Schramm(2) and adapted to include a cycling reaction to quantitate NAD+ production colorimetrically. The assay follows the NAMPT-catalyzed production of NMN from substrates NAM and PRPP by coupling NMN formation to the NMNAT reaction, which produces NAD+ from NMN and ATP. The NAD+ is then cycled by alcohol dehydrogenase (ADH) and diaphorase to continuously produce WST-1 formazan, which can be detected at 450 nm. Assays are performed at 25°C in clear 384-well plates, with a final assay volume of 30 uL, and contain the following: 50 mM HEPES, pH 7.5, 5 mM $MgCl_2$, 50 mM NaCl, 0.01% Triton-X 100, 2.5 mM ATP, 40 uM PRPP, 30 uM uM NAM, 1.5 uL WST-1 (Roche Cell Proliferation Reagent), 1U/mL ADH, 0.083 U/mL diaphorase, 1.5% ethanol, 1% DMSO, 30 nM NAMPT, and 7.4 nM purified human NMNAT1. All assay reagents were acquired from Sigma-Aldrich unless otherwise specified. N-terminal $His_6$-NMNAT1 was overexpressed and purified as detailed for NAMPT from an expression vector obtained from Genscript. Following assay assembly, well signals were measured continuously at 450 nm on a Tecan Infinite M200 plate reader for 1h with intermittent shaking. Slopes of the linear portions of the reaction progress curves were recorded and corrected for background by subtracting the average slope of control wells containing NAMPT inhibitor FK866.

A discontinuous version of the assay was used to examine the influence of N-PAMs on catalytic parameters with respect to ATP, PRPP, and NAM under steady state conditions. Briefly, 30 nM NAMPT was incubated with 2.5 mM ATP, 40 uM PRPP, and 30 uM NAM for a fixed period of time. Two substrate concentrations were held constant, while one was varied. The reaction was quenched with FK866, and cycling components (ADH, diaphorase, ethanol, WST-1) were added to measure NAD+. NAD+ concentrations were determined from a standard curve,

*NAMPT high-throughput screen.* The continuous primary assay described above was used for high-thoroughput screening of compound libraries. Pilot experiments demonstrated low variability of the signal across the 384-well plate with an excellent Z-factor (0.82). Using this HTS assay, we screened: (1) a 10,000 compound subset of the ChemDiv SMART™ Library and a 10,000 compound subset of the ChemDiv PPI Library, both chosen to balance the diversity of chemical scaffolds with favorable characteristics for future derivatization and to remove PAINs (pan-assay interference compounds; including oxidants, electrophiles, and undesirable functional groups (3, 4)); in addition to (2) 2,000 compounds from the Microsource Spectrum library. Inhibition of NMNAT by active compounds was ruled out by testing in the same assay with NAM replaced by NMN. Actives were further triaged to obtain validated hits by retesting repurchased or resynthesized compounds in triplicate at 8 concentrations and by use of orthogonal assays described below.

*Orthogonal NAMPT assay: NMN trapping.* The method of Zhang et al.(5) was modified and miniaturized to establish an orthogonal assay for NAMPT activity. The assay uses the trapping of NMN by acetophenone in basic solution followed by acidification to form a fluorescent adduct (380nm excitation, 450nm emission). Reactions were performed at RT for 60 min in 50mM Hepes (pH 7.5), 5mM MgCl2, 50mM NaCl, 30 μM NAMPT, 2.5 mM ATP, 30 μM NAM, 40 μM PRPP and test compound in 1% DMSO/buffer. After 60 min, to the reaction is added of 6.7 μL 20% acetophenone in DMSO and 6.7 μL 2M KOH then incubated on ice for 10 min. The reaction is quenched with 29 μL 88% formic acid, incubated for 15 min at 37 °C, and read immediately. Fluorescense was measured with a Clariostar Reader (excitation wavelength, 380 nm; emission wavelength, 450 nm). The NAMPT inhibitor FK866 was used to validate the assay. This is an endpoint assay in which NMN accumulates, in contrast to the primary NAMPT assay in which NMN is immediately captured by NMNAT. Since NAMPT is subject to feedback inhibition by substrates and products (including NMN and $NAD^+$), the orthogonal assay is not expected to yield identical data to the primary assay. The orthogonal enzyme activity assay was also used to study the ATP-independent interconversion of NAM with NMN and the shift in equilibrium on addition of ATP (2 mM): PRPP 100uM, PPi 100uM, NAMPT 40nM, NAM 3uM, NMN 3uM, SBI 5uM, NP-A1S 20uM (**Figs 4A, S10**).

*ATPase activity: non-stoichiometric turnover of ATP***.** Assay conditions to measure the ATPase activity of NAMPT were based on HPLC-based measurements performed by Burgos and Schramm .(2) The adapted assay uses the Promega ADP-Glo Kinase Assay to measure ADP production by NAMPT in the presence of various substrates and effectors. Assays were performed at room temperature and included 25 mM HEPES, pH 7.5, 50 mM NaCl, 5 mM $MgCl_2$, and 0.5 mM ATP. Reactions were initiated with the addition of 1.5 uM NAMPT, and aliquots were removed and quenched according to the manufactures protocol at various time points. Following a 30-minute incubation with ADP detection reagent, luminescence was measured on a Tecan Infinite M200 microplate reader. A 0-200 uM ADP standard curve was included for each experiment.

*NAMPT fluorescence polarization (FP) displacement assay*. All FP measurements were performed at room temperature in PBS buffer containing 0.01% Triton-X 100 and 1% DMSO in black 384-well plates. Measurements were performed on a Tecan F200 Pro plate reader fitted with polarized 485(20) nm emission filters and 535(25) nm emission filters. Initial titration of 20 nM FP probe with varying concentrations of NAMPT protein was performed to estimate a $K_d$ value for the probe (~750 nM), which defined the NAMPT concentration for all subsequent experiments. Anisotropy data from equilibrium competition binding experiments were fit to a four-parameter dose response equation in GraphPad prism. All FP experiments included the following control sets: unbound probe (probe alone), bound probe (probe + NAMPT), and background signal (NAMPT alone).

*Biophysical assay of ligand binding.* Microscale thermophoresis (MST) was used as an orthogonal biophysical assay, replicating the high affinity binding of a NAMPT inhibitor (NAMPTi) and the *R*-isomer of the N-PAM. MST experiments were performed on a Monolith NT.115pico (NanoTemper Technologies). Binding of 10 nM NAMPT (His-tag labeled) in PBS-P buffer to various concentrations of NP-A1-R, quercitrin, or NAMPT inhibitor was determined at 23.5°C, using Premium Coated Capillaries (NanoTemper Technologies), with 20% excitation power and 40% MST power.

### CELL BIOLOGY

*Control of cellular $NAD^+$ levels in THP-1 cell cultures.* Using FK-866 and NMN as negative and positive controls, respectively, we explored a number of cell lines to select a highly reproducible model system with good dynamic range. We optimized a commercial NADglo assay in the THP-1 human leukemic monocyte cell line in 96 well plates, measuring $NAD^+$ after incubation with test compounds for 24 h. The human monocytic leukemia cell line (THP-1) was obtained from the American Type Culture Collection and were maintained in RPMI 1640 medium (ATCC) supplemented with 100 units of penicillin, 100 µg/ml streptomycin, and 10% heat inactive fetal bovine serum. All cells were grown at 37 °C, under 5% $CO_2$ in a humidified incubator. Low passage THP-1 cells (37,500 cells/well) were seeded in 96-well plates and incubated at 37 °C and 5% $CO_2$ for 1 1/2 h prior to a 24 h treatment. All compounds were dissolved in DMSO, and final DMSO concentrations never exceeded 1%. The $NAD^+$ levels in the cells are measured using the NAD/NADH-Glo™ assay (Promega). The assays were performed in triplicate for each concentration and the $IC_{50}$ values were determined from non-linear regression analysis of the dose-response curve generated in GraphPad Prism 9.

### STRUCTURAL BIOLOGY

*Protein crystallography.* Crystals of all NAMPT complexes were grown by hanging drop vapor diffusion at 16°C. Prior to crystallization, 11 mg/mL NAMPT protein was incubated with excess molar concentrations of appropriate ligands for 30 min on ice. Crystals of the complexes were grown by mixing 1-2 uL of NAMPT complex with 2 uL of reservoir solution containing 0.1 M Tris-HCl, pH 8, 0.1 M KCl, and 24-28% PEG 2000 MME *or* 0.1 M Tris-HCl, pH 8.5, 0.2 M NaCl, 20% glycerol and 13-18% PEG 3350. Crystals grew overnight from the PEG 2000 MME conditions and were used to seed drops of NAMPT complex equilibrating against the PEG 3350 conditions.

*X-ray data collection and structure refinement.* The glycerol present in the crystallization solution was sufficient to cryo-protect crystals, which were flash-cooled in liquid nitrogen. Data were collected at the Life Sciences Collaborative Access Team beamlines 21-ID-F and 21-ID-G at the Advanced Photon Source, Argonne National Laboratory. Data indexing, integration, and scaling were performed using XDS(6) or Xia(7) and DIALS (8). Phases were determined by molecular replacement using Molrep(9) and a NAMPT-NAM co-crystal structure (PDB entry: 2E5D) as search model. Rigid body refinement followed by iterative rounds of restrained refinement and model building were performed with Phenix(10, 11) and Coot(12). The coordinates and structure factors of NAMPT complexes have been deposited with PDB accession codes as follows:

NAM + NP-A1-R (8DSC), NAM + NP-A1-S (8DSD), NAM + querictrin (8DSE), NAM (8DSI), quercitrin + AMPcP (8DSH), ZN-2-43S (8DTJ), ZN-4-3 (8DSM).

Final coordinates of 8DSC, 8DSD, and 8DSE each contain an ensemble of two states, representing the NAM-only state, in which waters occupy the activator site, and NAM-activator state, in which the activator displaces the waters. Water coordinates were obtained from the NAM-only structure (8DSI). Occupancy refinement for each state was performed by Phenix (10, 11) , as outlined by Pearce *et al.*(13)

This research used resources of the Advanced Photon Source, a U.S. Department of Energy (DOE) Office of Science User Facility operated for the DOE Office of Science by Argonne National Laboratory under Contract No. DE-AC02-06CH11357. Use of the LS-CAT Sector 21 was supported by the Michigan Economic Development Corporation and the Michigan Technology Tri-Corridor (Grant 085P1000817).

Normal mode and overlap analysis were performed using Bio3D version 2.3.0. Normal modes were calculated from alpha-carbon coordinates from crystal structures of apo NAMPT (pdb id: 2e5b) and various NAMPT complexes.(14) Trajectories of the first six nontrivial modes, which were shared by all complexes, were visualized in Pymol. The conformational difference vectors calculated from the transition from the apo state to the complexed states were used to identify which modes contribute to each large-scale motion.

**CHEMISTRY**

**General Chemical Experimental Information** Unless otherwise specified, reactions were performed under an inert atmosphere of argon and monitored by thin-layer chromatography (TLC) and/or LCMS. All reagents were purchased from commercial suppliers and used as provided. Synthetic intermediates were purified using CombiFlash chromatography system on 230−400 mesh silica gel. $^{1}H$ and $^{13}C$ NMR spectra were obtained using Bruker DPX-400, AVANCE-400 or AVANCE NEO-500 spectrometer at 400, 500 MHz and 100, 125 MHz, respectively. NMR chemical shifts were described in δ (ppm) using residual solvent peaks as standard (Chloroform-*d*, 7.26 ppm ($^{1}H$), 77.16 ppm ($^{13}C$); Methanol-*d4*, 3.31 ppm ($^{1}H$), 49.00 ppm ($^{13}C$); DMSO-*d6*, 2.50 ppm ($^{1}H$), 39.52 ppm ($^{13}C$)). Data were reported in a format as follows: chemical shift, multiplicity (s = singlet, d = doublet, dd = doublet of doublet, t = triplet, q = quartet, br = broad, m = multiplet, abq = ab quartet), number of protons, and coupling constants. High resolution mass spectral data were measured in-house using a Shimadzu IT-TOF LC/MS for final compounds. All compounds submitted for biological testing were confirmed to be ≥ 95% pure by analytical HPLC. Synthetic methods, spectral data, and HRMS for novel compounds are described in detail below.

**General synthetic routes for compounds NP-A1S, NP-A1R and ZN-2-43S**

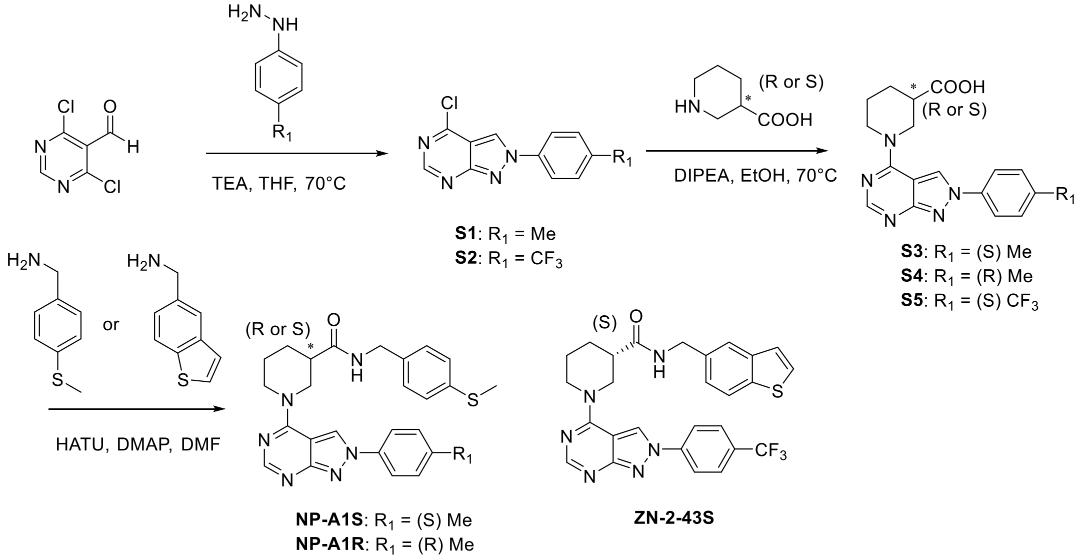

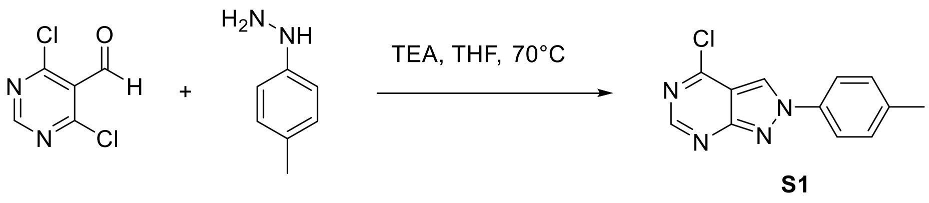

**4-chloro-2-(p-tolyl)-2H-pyrazolo[3,4-d]pyrimidine (S1)**. To a round bottom flask was added 4,6-dichloropyrimidine-5-carboxaldehyde (500 mg, 2.83 mmol), p-tolylhydrazine hydrochloride (449 mg, 2.83 mmol), and THF (20 mL). The reaction was allowed to stir, under a nitrogen atmosphere, at 70 °C for 1 hour. After removal of solvent, the residue was purified by silica gel column chromatography (Hexenes/EtOAc, 2:1) to provide the 4-chloro-2-(p-tolyl)-2H-pyrazolo[3,4-d]pyrimidine (291 mg, 42%) as a white solid: $^1$H NMR (400 MHz, Chloroform-*d*) δ 8.86 (s, 1H), 8.33 (s, 1H), 8.02 (d, *J* = 8.5 Hz, 2H), 7.36 (d, *J* = 8.2 Hz, 2H), 2.44 (s, 3H); LRMS (ESI) calcd for $C_{12}H_{10}ClN_4$ [M + H]$^+$ 245.06, found 245.00.

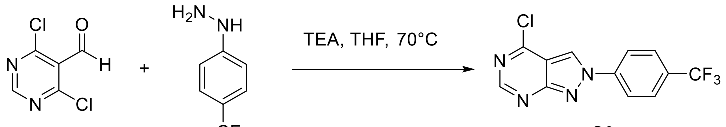

S2

**4-chloro-2-(4-(trifluoromethyl)phenyl)-2H-pyrazolo[3,4-d]pyrimidine (S2)**. To a round bottom flask was added 4,6-dichloropyrimidine-5-carboxaldehyde (500 mg, 2.83 mmol), (4-(trifluoromethyl)phenyl)hydrazine hydrochloride (600 mg, 2.83 mmol), and THF (20 mL). The reaction was allowed to stir, under a nitrogen atmosphere, at 70 °C for 1 hour. After removal of solvent, the residue was purified by silica gel column chromatography (Hexenes/EtOAc, 2:1) to provide the desired compound (293 mg, 35%) as a light brown solid: $^1$H NMR (400 MHz, Chloroform-*d*) δ 8.93 (s, 1H), 8.70 (s, 1H), 8.14 (d, *J* = 8.5 Hz, 2H), 7.87 (d, *J* = 8.5 Hz, 2H); LRMS (ESI) calcd for $C_{12}H_7ClF_3N_4$ [M + H]$^+$ 299.03, found 299.10.

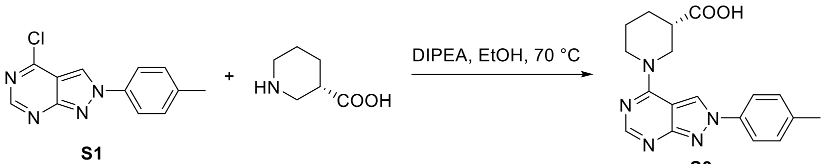

S1 S3

**(S)-1-(2-(p-tolyl)-2H-pyrazolo[3,4-d]pyrimidin-4-yl)piperidine-3-carboxylic acid (S3)**. To a solution of **S1** (47 mg, 0.19 mmol) and (S)-piperidine-3-carboxylic acid (29 mg, 0.23 mmol) in EtOH (1 mL), DIPEA (200 µL) was added. After stirring at 70 °C for 1 h and cooldown, the solvent was removed under vacuum. The residue was purified by Prep-HPLC to provide the desired compound (59 mg, yield 92%) as a light-yellow solid: $^1$H NMR (500 MHz, Methanol-*d*$_4$) δ 9.11 (s, 1H), 8.33 (s, 1H), 7.95 – 7.80 (m, 2H), 7.47 – 7.37 (m, 2H), 4.38 (dd, *J* = 13.4, 4.7 Hz, 1H), 3.84 – 3.60 (m, 2H), 2.76 – 2.68 (m, 1H), 2.45 (s, 3H), 2.23 – 2.15 (m, 1H), 2.01 – 1.91 (m, 2H), 1.77 – 1.67 (m, 1H), 1.38 – 1.23 (m, 1H); LRMS (ESI) calcd for $C_{18}H_{20}N_5O_2$ [M + H]$^+$ 338.16, found 338.12.

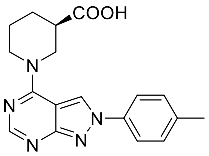

**S4**

**(R)-1-(2-(p-tolyl)-2H-pyrazolo[3,4-d]pyrimidin-4-yl)piperidine-3-carboxylic acid (S4)**. Light-yellow solid (yield 92%): $^1$H NMR (500 MHz, Methanol-*d*$_4$) δ 9.09 (s, 1H), 8.30 (s, 1H), 7.91 – 7.84 (m, 2H), 7.42 – 7.30 (m, 2H), 4.36 (dd, *J* = 11.4, 6.8 Hz, 1H), 3.76 – 3.65 (m, 2H), 2.73 – 2.66 (m, 1H), 2.43 (s, 3H), 2.19 – 2.12 (m, 1H), 2.00 – 1.90 (m, 2H), 1.76 – 1.66 (m, 1H), 1.33 – 1.25 (m, 1H); LRMS (ESI) calcd for $C_{18}H_{20}N_5O_2$ [M + H]$^+$ 338.16, found 338.11.

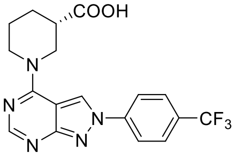

**S5**

**(S)-1-(2-(4-(trifluoromethyl)phenyl)-2H-pyrazolo[3,4-d]pyrimidin-4-yl)piperidine-3-carboxylic acid (S5)**. White solid (yield 93%); $^1$H NMR (400 MHz, Methanol-*d*$_4$) δ 9.20 (s, 1H), 8.28 (s, 1H), 8.19 (t, *J* = 7.5 Hz, 2H), 7.84 (d, *J* = 8.4 Hz, 2H), 4.51 – 4.14 (m, 2H), 3.84 – 3.58 (m, 2H), 2.79 – 2.63 (m, 1H), 2.23 – 2.11 (m, 1H), 2.00 – 1.86 (m, 2H), 1.78 – 1.64 (m, 1H); LRMS (ESI) calcd for $C_{18}H_{19}FN_5O_2$ [M + H]$^+$ 392.13, found 392.14.

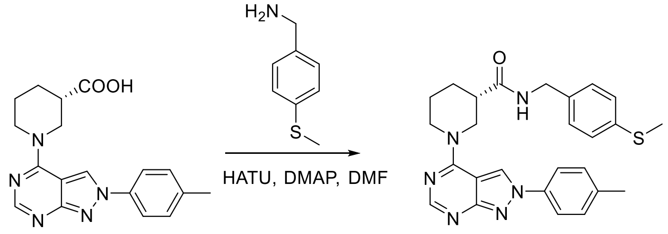

**S3** **NP-A1S**

**(S)-N-(4-(methylthio)benzyl)-1-(2-(p-tolyl)-2H-pyrazolo[3,4-d]pyrimidin-4-yl)piperidine-3-carboxamide (NP-A1S)**. **S3** (34 mg, 0.10 mmol), (4-(methylthio)phenyl)methanamine (30 mg, 0.2 mmol), HATU (57 mg, 0.15 mmol), and DMAP (37 mg, 0.3 mmol) were dissolved in dry DMF (1 mL) and stirred at room temperature overnight. The mixture was diluted with Ethyl Acetate and was then washed with saturated aq. $NaHCO_3$, water, and brine, respectively. The organic layer was dried over $Na_2SO_4$, filtered, and concentrated. The residue was purified by Prep-HPLC to provide the **NP-A1S** (41 mg, 87%) as a white solid: $[\alpha]_D^{25}$ = +94.5 (c 0.5, MeOH); $^1$H NMR (400 MHz, DMSO-*d*$_6$) δ 9.27 (s, 1H), 8.45 (t, *J* = 5.7 Hz, 1H), 8.27 (s, 1H), 7.97 (d, *J* = 8.1 Hz, 2H), 7.39 (d, *J* = 8.2 Hz, 2H), 7.26 – 7.04 (m, 4H), 5.02 – 4.43

(m, 2H), 4.31 – 4.17 (m, 2H), 3.56 – 3.14 (m, 2H), 2.47 – 2.46 (m, 1H), 2.44 (s, 3H), 2.39 (s, 3H), 1.98 – 1.77 (m, 3H), 1.61 – 1.46 (m, 1H). $^{13}$C NMR (100 MHz, DMSO-*d*$_6$) δ 172.90, 161.33, 157.90, 156.31, 138.15, 137.58, 136.75, 136.69, 130.44, 128.37, 126.57, 123.81, 120.54, 102.72, 47.19, 46.55, 42.64, 41.97, 28.10, 24.84, 21.03, 15.41; HRMS (ESI) calcd for $C_{26}H_{29}N_6OS$ [M+H]$^+$ 473.2118, found 473.2127.

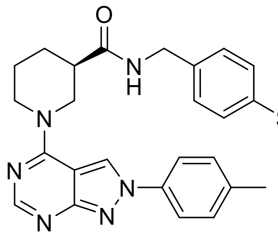

**NP-A1R**

**(R)-N-(4-(methylthio)benzyl)-1-(2-(p-tolyl)-2H-pyrazolo[3,4-d]pyrimidin-4-yl)piperidine-3-carboxamide (NP-A1R)**. White solid (yield 84%): $[\alpha]_D^{25}$ = -87.8 (c 0.7, MeOH); $^1$H NMR (400 MHz, DMSO-*d*$_6$) δ 9.27 (s, 1H), 8.46 (t, *J* = 6.0 Hz, 1H), 8.27 (s, 1H), 7.97 (d, *J* = 8.2 Hz, 2H), 7.39 (d, *J* = 8.2 Hz, 2H), 7.19 (s, 4H), 5.03 – 4.47 (m, 2H), 4.33 – 4.16 (m, 2H), 3.43 – 3.25 (m, 2H), 2.47 – 2.46 (m, 1H), 2.44 (s, 3H), 2.39 (s, 3H), 2.00 – 1.75 (m, 3H), 1.61 – 1.46 (m, 1H); $^{13}$C NMR (100 MHz, DMSO-*d*$_6$) δ 172.43, 160.86, 157.43, 155.83, 137.68, 137.11, 136.27, 136.22, 129.97, 127.90, 126.09, 123.34, 120.07, 102.24, 46.72, 45.32, 42.11, 41.50, 27.62, 24.35, 20.56, 14.93; HRMS (ESI) calcd for $C_{26}H_{29}N_6OS$ [M+H]$^+$ 473.2118, found 473.2125.

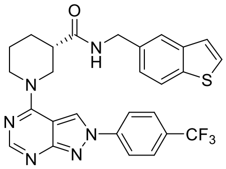

**ZN-2-43S**

**(S)-N-(benzo[b]thiophen-5-ylmethyl)-1-(2-(4-(trifluoromethyl)phenyl)-2H-pyrazolo[3,4-d]pyrimidin-4-yl)piperidine-3-carboxamide (ZN-2-43S)**. Light brown solid (yield 87%): $[\alpha]_D^{25}$ = +83.4 (c 0.5, MeOH); $^1$H NMR (400 MHz, DMSO-*d*$_6$) δ 9.48 (s, 1H), 8.54 (t, *J* = 5.5 Hz, 1H), 8.34 (d, *J* = 8.3 Hz, 2H), 8.30 (s, 1H), 8.01 – 7.88 (m, 3H), 7.79 – 7.70 (m, 2H), 7.48 – 7.37 (m, 1H), 7.28 (d, *J* = 8.3 Hz, 1H), 5.13 – 4.52 (m, 2H), 4.41 (d, *J* = 5.7 Hz, 2H), 3.48 – 3.37 (m, 2H), 2.64 – 2.54 (m, 1H), 2.08 – 1.77 (m, 3H), 1.56 (d, *J* = 12.1 Hz, 1H); $^{13}$C NMR (100 MHz, DMSO-*d*$_6$) δ 172.39, 161.12, 157.51, 156.50, 142.13, 139.52, 137.63, 135.71, δ 128.01 (q, *J* = 32.5 Hz), 127.75, 126.88, 124.62, 124.07, 123.97 (q, *J* = 271.5 Hz), 123.79, 122.43, 122.08, 120.62, 102.87, 47.00, 46.39, 42.24, 42.03, 27.63, 24.66; HRMS (ESI) calcd for $C_{27}H_{24}F_3N_6OS$ [M+H]$^+$ 537.1679, found 537.1686.

**Synthetic route for compound ZN-2-102**

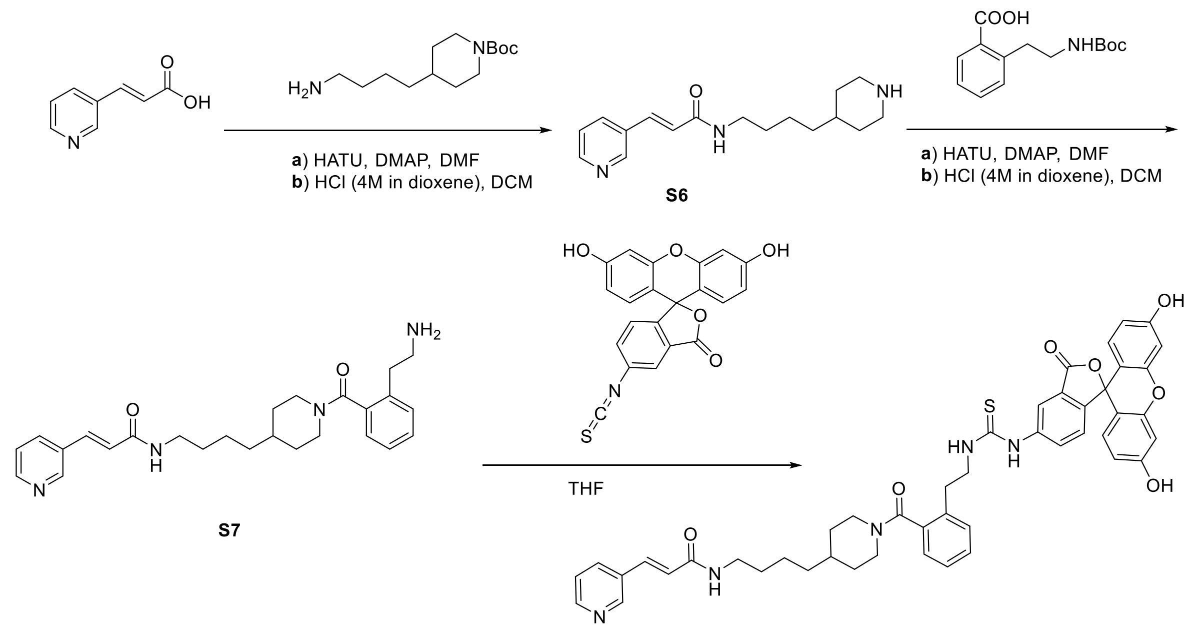

**ZN-2-102**

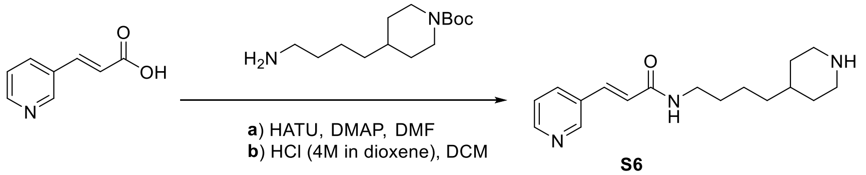

**S6**

**(E)-N-(4-(piperidin-4-yl)butyl)-3-(pyridin-3-yl)acrylamide (S6)**. (E)-3-(pyridin-3-yl)acrylic acid (150 mg, 1.0 mmol), tert-butyl 4-(4-aminobutyl)piperidine-1-carboxylate (256 mg, 1.0 mmol), HATU (456 mg, 1.2 mmol) and DMAP (366 mg, 3.0 mmol) were dissolved in dry DMF (4 mL) and stirred at room temperature overnight. After purification by Prep-HPLC, the product was dissolved in DCM (5 mL) at 0 °C. HCl (4M in dioxane, 1.3 mL) was added slowly into the system and warmed up to room temperature. After stirring for another 0.5 h, the reaction was dried under vacuum to afford the compound **S6** (241 mg, yield 84% for 2 steps) as a white solid: $^1$H NMR (500 MHz, Methanol-$d_4$) δ 8.80 – 8.47 (m, 3H), 8.05 (d, $J$ = 7.7 Hz, 1H), 7.59 – 7.44 (m, 2H), 6.73 (d, $J$ = 15.7 Hz, 1H), 3.46 – 3.34 (m, 4H), 3.03 – 2.89 (m, 2H), 1.95 (d, $J$ = 12.8 Hz, 2H), 1.67 – 1.23 (m, 9H); LRMS (ESI) calcd for $C_{17}H_{26}N_3O$ [M+H]$^+$ 288.21, found 288.17.

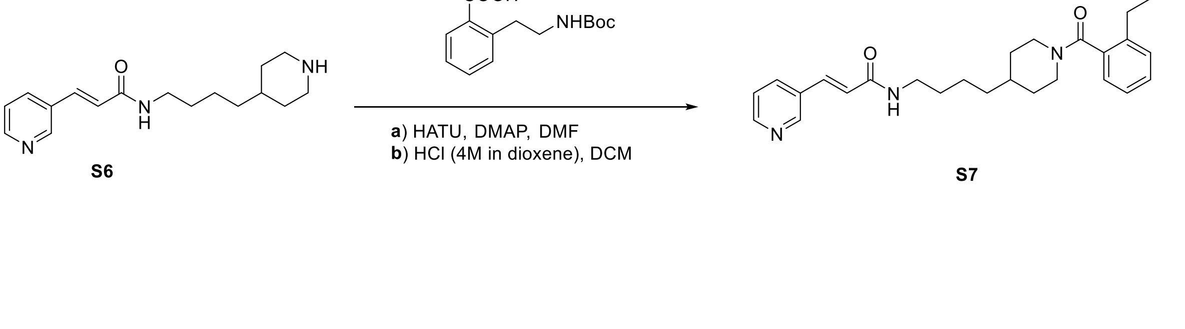

**(E)-N-(4-(1-(2-(2-aminoethyl)benzoyl)piperidin-4-yl)butyl)-3-(pyridin-3-yl)acrylamide (S7)**. **S6** (150 mg, 0.52 mmol), 2-(2-((tert-butoxycarbonyl)amino)ethyl)benzoic acid (160 mg, 0.63 mmol), HATU (339 mg, 0.78 mmol) and DMAP (191 mg, 1.56 mmol) were dissolved in dry DMF (4 mL) and stirred at room temperature overnight. After purification by Prep-HPLC, the product was dissolved in DCM (4 mL) at 0 °C. HCl (4M in dioxane, 650 μL) was added slowly into the system and warmed up to room temperature. After stirring for another 0.5 h, the reaction was dried under vacuum to afford the compound **S7** (192 mg, yield 85% for 2 steps) as a white solid: $^{1}$H NMR (500 MHz, Methanol-*d*$_4$) δ 8.71 (s, 1H), 8.52 (d, *J* = 4.9 Hz, 1H), 8.36 (s, 1H), 8.04 (d, *J* = 7.9 Hz, 1H), 7.54 (d, *J* = 15.9 Hz, 1H), 7.51 – 7.35 (m, 4H), 7.32 – 7.21 (m, 1H), 6.73 (d, *J* = 15.8 Hz, 1H), 4.73 – 4.65 (m, 1H), 3.53 – 3.40 (m, 1H), 3.35 – 3.31 (m, 2H), 3.23 – 2.74 (m, 6H), 1.93 – 1.86 (m, 1H), 1.73 – 1.53 (m, 4H), 1.47 – 1.15 (m, 6H); LRMS (ESI) calcd for $C_{26}H_{35}N_4O_2$ [M+H]$^+$ 435.28, found 435.30.

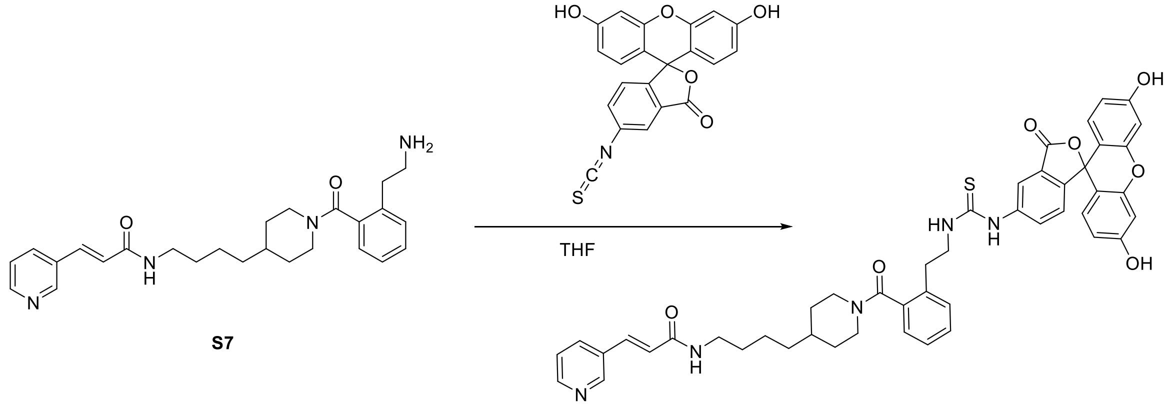

**(E)-N-(4-(1-(2-(2-(3-(3',6'-dihydroxy-3-oxo-3H-spiro[isobenzofuran-1,9'-xanthen]-5-yl)thioureido)ethyl)benzoyl)piperidin-4-yl)butyl)-3-(pyridin-3-yl)acrylamide (ZN-2-102)**. **S7** (10 mg, 0.023 mmol) and 3',6'-dihydroxy-5-isothiocyanato-3H-spiro[isobenzofuran-1,9'-xanthen]-3-one (13.4 mg, 0.035 mmol) were dissolved in THF (1 mL). The mixture was stirred at room temperature until the reaction finished. The reaction was purified by Prep-HPLC to provide the yellow solid **ZN-2-102** (12 mg, yield 63%): $^{1}$H NMR (500 MHz, Methanol-*d*$_4$) δ 8.71 – 8.65 (m, 1H), 8.50 (dt, *J* = 4.6, 1.9 Hz, 1H), 8.01 (d, *J* = 6.1 Hz, 2H), 7.66 – 7.38 (m, 5H), 7.30 (q, *J* = 8.3 Hz, 1H), 7.25 – 7.05 (m, 2H), 6.74 – 6.62 (m, 5H), 6.57 – 6.50 (m, 2H), 4.62 (d, *J* = 13.0 Hz, 1H), 3.95 – 3.76 (m, 2H), 3.55 – 3.43 (m, 1H), 3.31 – 3.26 (m, 2H), 3.08 (t, *J* = 12.9 Hz, 2H), 2.88 – 2.74 (m, 2H), 1.83 (d, *J* = 13.2 Hz, 1H), 1.64 (d, *J* = 13.3 Hz, 1H), 1.56 (q, *J* = 7.1 Hz, 3H), 1.43 – 1.08 (m, 6H); $^{13}$C NMR (125 MHz, MeOD) δ 171.08, 167.68, 161.41, 154.19, 150.66, 149.67, 137.36, 136.25, 132.91, 130.62, 130.30, 127.87, 127.27, 126.99, 125.53, 124.82, 113.60, 111.48, 103.54, 43.26, 40.54, 37.18, 37.03, 33.13, 30.54, 24.98; HRMS (ESI) calcd for $C_{47}H_{46}N_5O_7S$ [M+H]$^+$ 824.3112, found 824.3120.

### Synthetic route for compound ZN-4-3

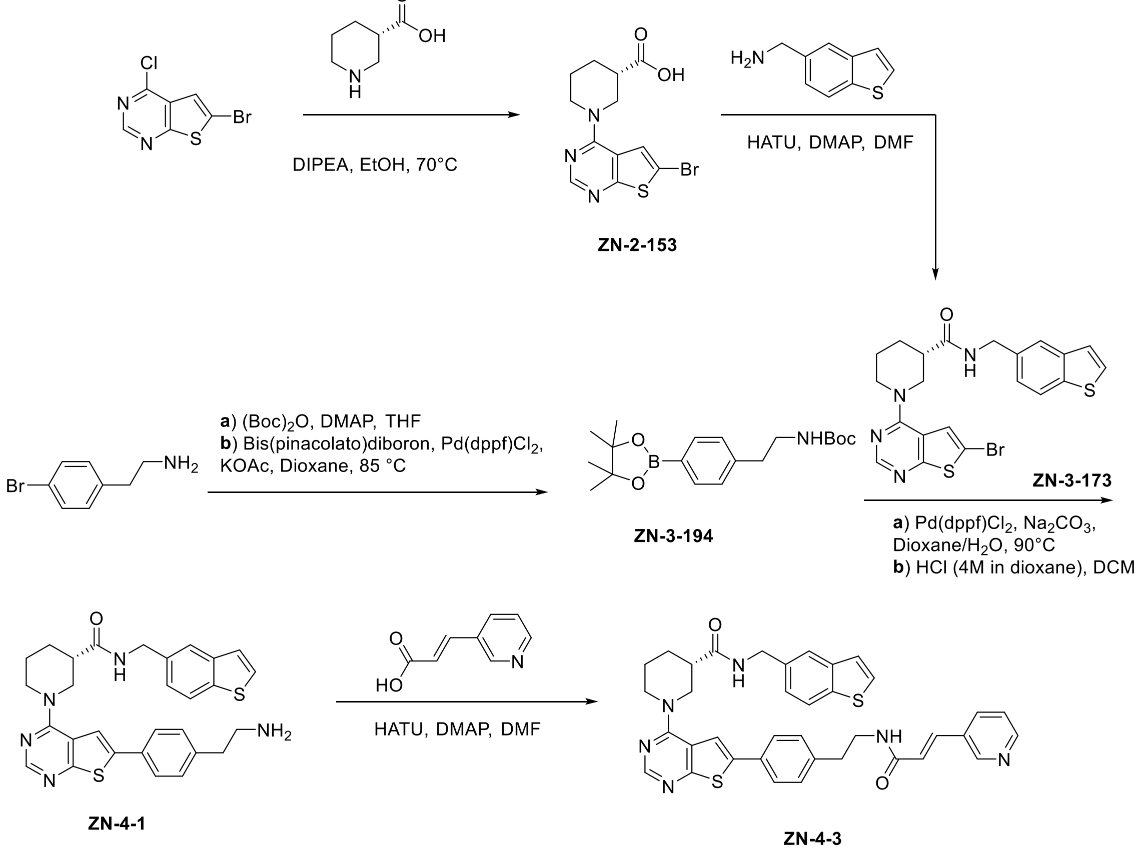

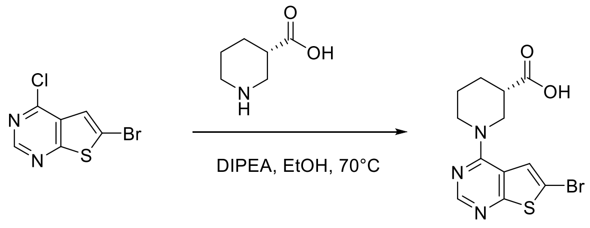

**S8**

**(S)-1-(6-bromothieno[2,3-d]pyrimidin-4-yl)piperidine-3-carboxylic acid (S8)**. To a solution of 6-bromo-4-chlorothieno[2,3-d]pyrimidine (250 mg, 1.0 mmol) and (S)-piperidine-3-carboxylic acid (194 mg, 1.5 mmol) in EtOH (5 mL), DIPEA (1 mL) was added. After stirring at 70 °C for 1 h and cooldown, the solvent was removed under vacuum. The residue was purified by Prep-HPLC to provide the **S8** (311 mg, yield 91%) as a white solid: $^1$H NMR (500 MHz, Methanol-$d_4$) δ 8.32 (s, 1H), 7.68 (s, 1H), 4.51 (dd, *J* = 13.6, 3.7 Hz, 1H), 4.26 (dd, *J* = 13.5, 4.4 Hz, 1H), 3.53 (dd, *J* = 13.4, 9.6 Hz, 1H), 3.43

(ddd, *J* = 13.5, 10.7, 3.2 Hz, 1H), 2.63 – 2.57 (m, 1H), 2.16 – 2.09 (m, 1H), 1.91 – 1.82 (m, 2H), 1.70 – 1.61 (m, 1H); LRMS (ESI) calcd for $C_{12}H_{13}BrN_3O_2S$ $[M+H]^+$ 341.99, found 341.82.

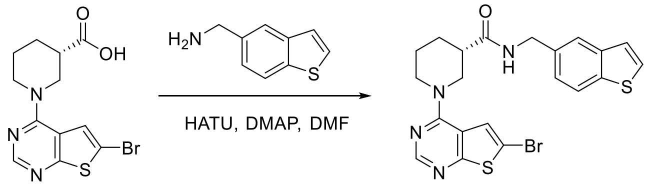

S8 S9

**(S)-N-(benzo[b]thiophen-5-ylmethyl)-1-(6-bromothieno[2,3-d]pyrimidin-4-yl)piperidine-3-carboxamide (S9)**. **S8** (200 mg, 0.58 mmol), benzo[b]thiophen-5-ylmethanamine (142 mg, 0.87 mmol), HATU (331 mg, 0.87 mmol), and DMAP (213 mg, 1.74 mmol) were dissolved in dry DMF (4 mL) and stirred at room temperature overnight. The mixture was purified by Prep-HPLC to provide the **S9** (237 mg, 84%) as a white solid: $^1$H NMR (500 MHz, Methanol-*d*$_4$) δ 8.28 (s, 1H), 7.84 (d, *J* = 8.3 Hz, 1H), 7.75 (s, 1H), 7.61 – 7.53 (m, 2H), 7.30 (dd, *J* = 24.5, 6.9 Hz, 2H), 4.51 (dt, *J* = 27.6, 14.3 Hz, 4H), 3.56 – 3.48 (m, 1H), 3.38 (t, *J* = 12.7 Hz, 1H), 2.66 – 2.59 (m, 1H), 2.11 – 1.84 (m, 3H), 1.72 – 1.61 (m, 1H); LRMS (ESI) calcd for $C_{21}H_{20}BrN_4OS_2$ $[M+H]^+$ 487.03, found 487.10.

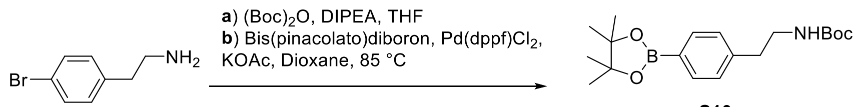

S10

**tert-butyl (4-(4,4,5,5-tetramethyl-1,3,2-dioxaborolan-2-yl)phenethyl)carbamate (S10)**. 2-(4-bromophenyl)ethan-1-amine (74 mg, 0.37 mmol), $(Boc)_2O$ (100 μL, 0.43 mmol) and DIPEA (322 μL, 1.85 mmol) were dissolved in THF (4 mL) and stirred at room temperature for 2 h. After being quenched with methanol, the mixture was purified by silica gel column chromatography (Hexenes/EtOAc, 5:1) to provide the intermediate. A flask fitted with a rubber septum was charged with the intermediate, Bis(pinacolato)diboron (109 mg, 0.43 mmol), Pd(dppf)$Cl_2$ (29 mg, 0.04 mmol), KOAc (73 mg, 0.74 mmol), Dioxane (4 mL) and then purged with argon. After stirring at 85 $^o$C overnight, the reaction mixture was then cooled to room temperature, diluted with ethyl acetate (20 mL), filtered through celite and concentrated in vacuo. The purification by Prep-HPLC afforded **S10** (60 mg, yield 47% over 2 steps) as a white solid: $^1$H NMR (500 MHz, Chloroform-*d*) δ 7.78 – 7.73 (m, 2H), 7.20 (d, *J* = 7.5 Hz, 2H), 3.37 (q, *J* = 6.8 Hz, 2H), 2.81 (t, *J* = 7.0 Hz, 2H), 1.43 (s, 9H), 1.34 (s, 12H); LRMS (ESI) calcd for $C_{19}H_{31}BrNO_4$ $[M+H]^+$ 348.23, found 348.33.

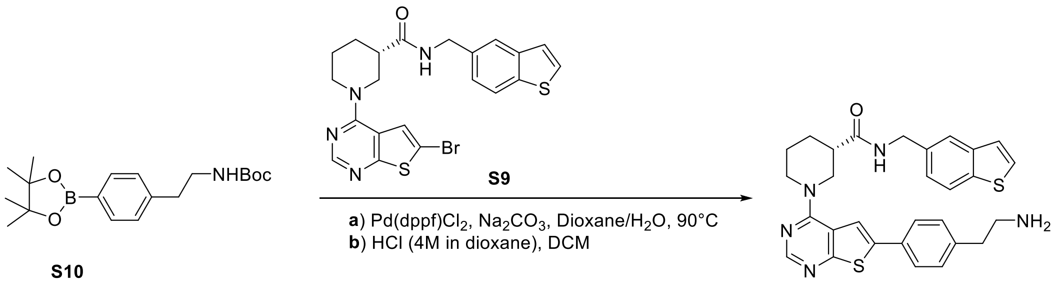

S10

S11

**(S)-1-(6-(4-(2-aminoethyl)phenyl)thieno[2,3-d]pyrimidin-4-yl)-N-(benzo[b]thiophen-5-ylmethyl)piperidine-3-carboxamide (S11)**. A flask fitted with a rubber septum was charged with **S9** (49 mg, 0.10 mmol), **S10** (52 mg, 0.15 mmol), Pd(dppf)$Cl_2$ (8 mg, 0.01 mmol), $Na_2CO_3$ (32 mg, 0.3 mmol), Dixoane/$H_2O$ (4 mL/ 1 mL) and then purged with argon. The mixture was stirred at 90 $^{o}$C overnight. The reaction mixture was then cooled to room temperature, diluted with ethyl acetate (20 mL), filtered through celite and concentrated in vacuo. After purification by Prep-HPLC, the product was subjected to N-Boc deprotection procedure with HCl (4M in dioxane, 100 µL) and DCM (2 mL). The purification by Prep-HPLC afforded the product **S11** (41 mg, yield 77% over 2 steps) as a light yellow solid: $^{1}$H NMR (500 MHz, Methanol-*d*$_4$) δ 8.55 (s, 1H), 8.30 (s, 1H), 7.80 (d, *J* = 8.3 Hz, 1H), 7.75 (s, 1H), 7.71 (s, 1H), 7.70 – 7.64 (m, 2H), 7.54 (d, *J* = 5.4 Hz, 1H), 7.28 (td, *J* = 8.0, 3.5 Hz, 4H), 4.66 – 4.50 (m, 4H), 4.48 – 4.40 (m, 1H), 3.59 (dd, *J* = 13.4, 10.2 Hz, 1H), 3.46 – 3.37 (m, 1H), 3.18 – 3.12 (m, 1H), 2.96 (t, *J* = 7.8 Hz, 2H), 2.68 (tt, *J* = 10.4, 4.0 Hz, 1H), 2.08 (dd, *J* = 13.2, 4.3 Hz, 1H), 2.04 – 1.88 (m, 2H), 1.76 – 1.65 (m, 1H); LRMS (ESI) calcd for $C_{29}H_{30}N_5O_4S_2$ $[M+H]^+$ 528.19, found 528.17.

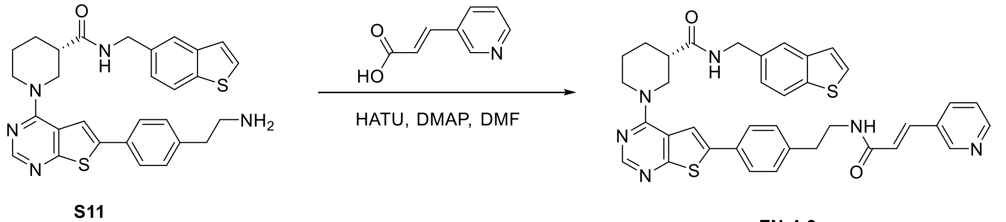

S11

ZN-4-3

**(S)-1-(6-(4-(2-aminoethyl)phenyl)thieno[2,3-d]pyrimidin-4-yl)-N-(benzo[b]thiophen-5-ylmethyl)piperidine-3-carboxamide (ZN-4-3)**. **S11** (20 mg, 0.038 mmol), (E)-3-(pyridin-3-yl)acrylic acid (11 mg, 0.076 mmol), HATU (29 mg, 0.076 mmol), and DMAP (14 mg, 0.114 mmol) were dissolved in dry DMF (1 mL) and stirred at room temperature overnight. The mixture was purified by Prep-HPLC to provide the **ZN-4-3** (22 mg, 88%) as a white solid: $^{1}$H NMR (500 MHz, Chloroform-*d*) δ 8.74 (s, 1H), 8.56 (s, 1H), 8.20 (s, 1H), 8.08 (s, 1H), 7.79 (dd, *J* = 8.3, 2.7 Hz, 2H), 7.70 (d, *J* = 1.6 Hz, 1H), 7.62 (d, *J* = 15.7 Hz, 1H), 7.59 – 7.57 (m, 2H), 7.48 (s, 1H), 7.45 (d, *J* = 5.5 Hz, 1H), 7.33 (dd, *J*

= 8.0, 4.9 Hz, 1H), 7.29 – 7.21 (m, 3H), 6.42 (d, *J* = 15.6 Hz, 1H), 5.78 (t, *J* = 5.9 Hz, 1H), 4.63 – 4.46 (m, 2H), 4.22 – 4.05 (m, 3H), 3.73 – 3.65 (m, 3H), 2.94 (t, *J* = 6.9 Hz, 2H), 2.66 (tt, *J* = 7.9, 4.0 Hz, 1H), 2.23 (dtd, *J* = 12.7, 8.6, 4.0 Hz, 1H), 2.03 – 1.96 (m, 1H), 1.83 – 1.66 (m, 2H); LRMS (ESI) calcd for $C_{37}H_{35}N_6O_2S_2$ $[M+H]^+$ 659.23, found 659.22.

### TABLES AND FIGURES

**Table S1.** Kinetic parameters for NAMPT activity dependence on substrates

| | ATP | | | PRPP | | | NAM | | | | |
|---|---|---|---|---|---|---|---|---|---|---|---|
| | $V_{max}$, min | $K_M$, mM | V/K | $V_{max}$, min | $K_M$, µM | V/K | $V_{max}$, min | $K_M$, µM | $V/K_M$ | $K_I$, µM | $V/K_I$ |
| **control** | 0.33±0.03 | 1.28±0.22 | 0.26±0.07 | 0.690±0.018 | 0.272±0.030 | 2.54±0.35 | 0.805±0.061 | 0.145±0.023 | 5.55±1.30 | 12±3 | 0.067±0.022 |
| **NP-A1R** | 0.98±0.05 | 1.51±0.15 | 0.65±0.10 | 0.966±0.023 | 0.356±0.035 | 2.72±0.33 | 0.717±0.016 | 0.149±0.082 | 4.81_2.76 | 289±43 | 0.0025±0.0004 |
| **NP-A1S** | 1.94±0.06 | 0.80±0.06 | 2.44±0.25 | 2.236±0.045 | 0.878±0.060 | 2.55±0.23 | 1.747±0.075 | 0.384±0.062 | 4.55±0.93 | 121±23 | 0.0144±0.0034 |
| **Quercitrin** | 1.24±0.07 | 0.30±0.05 | 4.20±0.94 | 1.730±0.040 | 1.001±0.073 | 1.73±0.17 | 1.637±0.055 | 0.483±0.050 | 3.39±0.46 | 40±4 | 0.041±0.005 |

Data obtained using primary coupled enzyme assay showing mean and SD from triplicate measurements

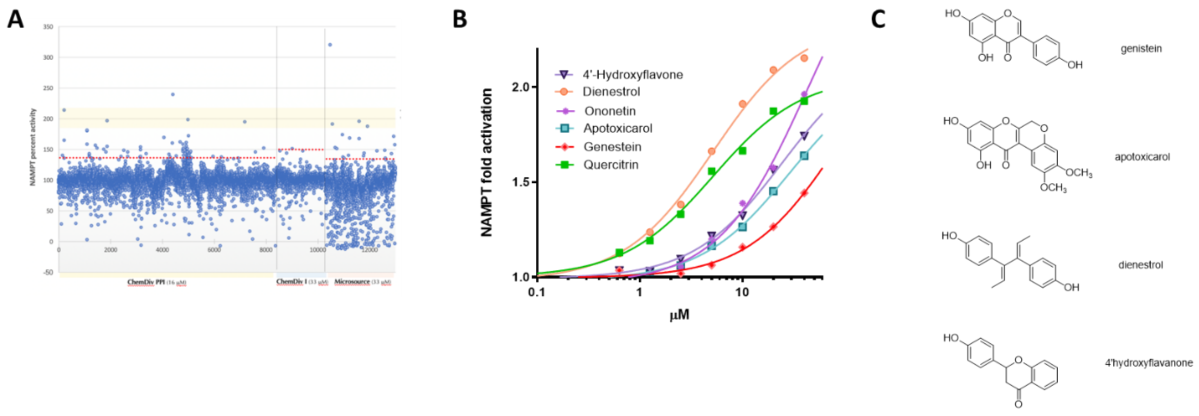

**Fig. S1.** A) Representative raw screening data from HTS (primary, coupled enzyme activity assay) of ChemDiv and Spectrum libraries. B) Concentration-response for hits from the Spectrum library with structures shown (C).

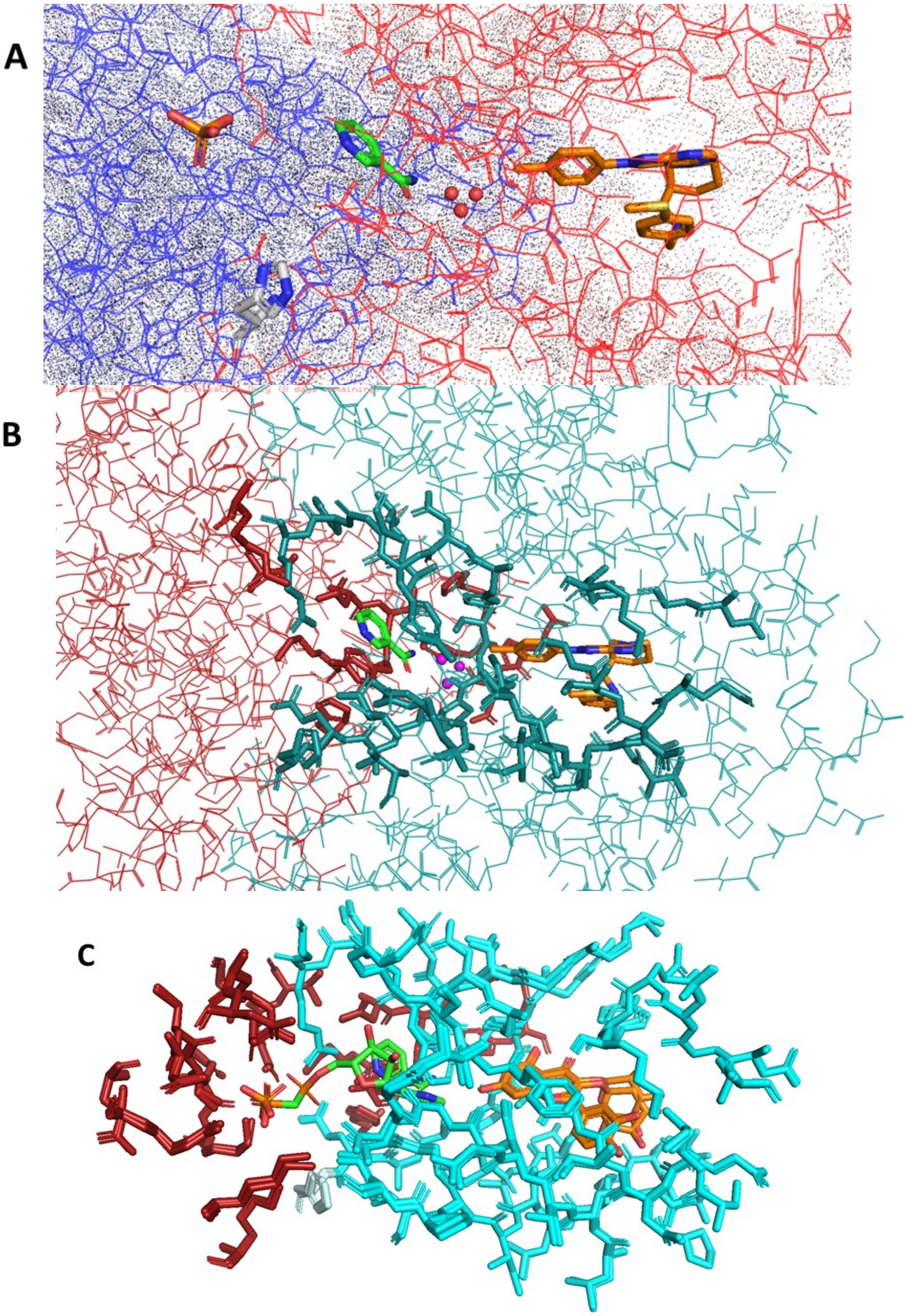

**Figure S2.** (A) Colorization of NAMPT monomers in blue and red emphasizes the location of the active site, nucleobase pocket and rear channel at the dimer interface: phosphate, His-247, water H-bond network and NP-A1S are highlighted. (B) Structure of NP-A1R (gold) bound to NAMPT in the presence of NAM (green) showing 3 waters (magenta) and dimer interface with residues lining rear channel and nucleobase emphasized. (C) Superposition of the two co-crystal structures of NAMPT-bound quercitrin (orange) showing NAM and the ADP analogue (AMPCP) bound to the nucleobase binding site. The terminal phosphonate of AMPCP superposes with a phosphate in the NAM-bound structure. Residues proximal to the rear channel and active site are shown with the two monomers in red or cyan. His-247 is shown in pale cyan.

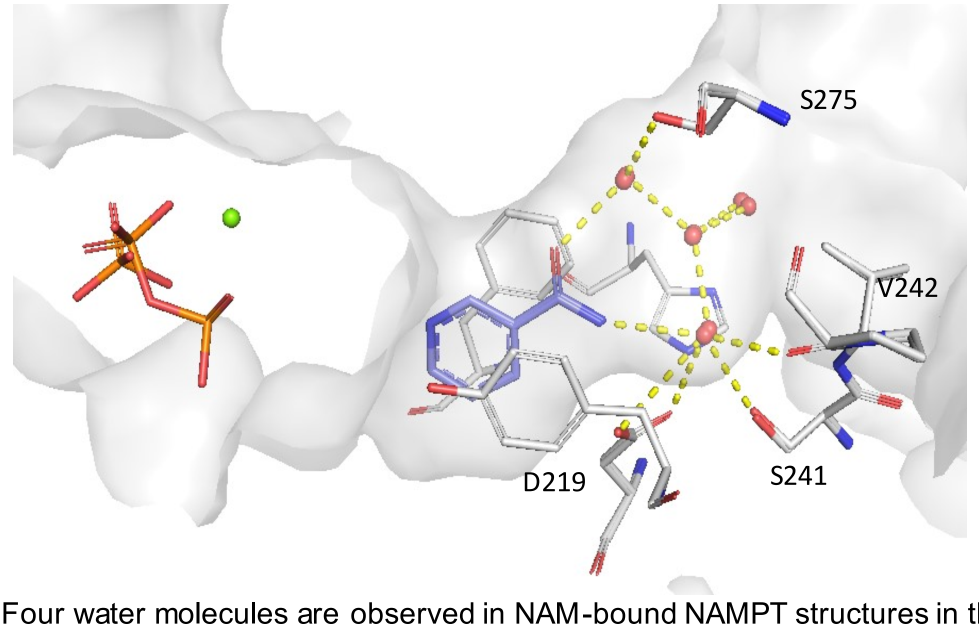

**Figure S3.** Four water molecules are observed in NAM-bound NAMPT structures in the absence of a rear channel ligand, with three water molecules superposing with the three water H-bonding network observed in N-PAM bound co-crystal structures.

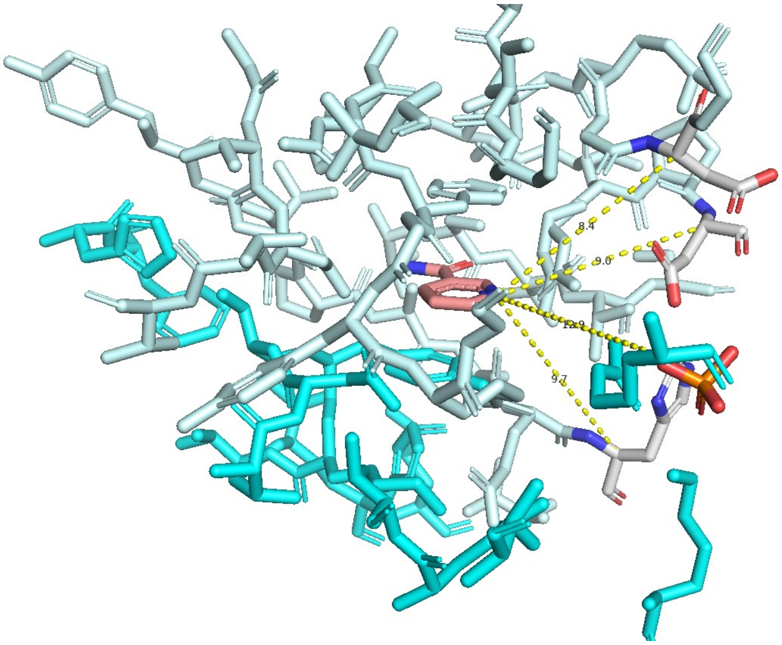

**Figure S4.** Structure of NAM (salmon) bound to the NAMPT nucleobase site showing the triangulation of the NAM nitrogen with α-carbons of active site residues that was used to measure the effect of ligand binding to the rear channel on the position of NAM. Binding of ligands to the rear channel had no effect on the four distance measurements shown. The two NAMPT monomers are coloured cyan and pale cyan.

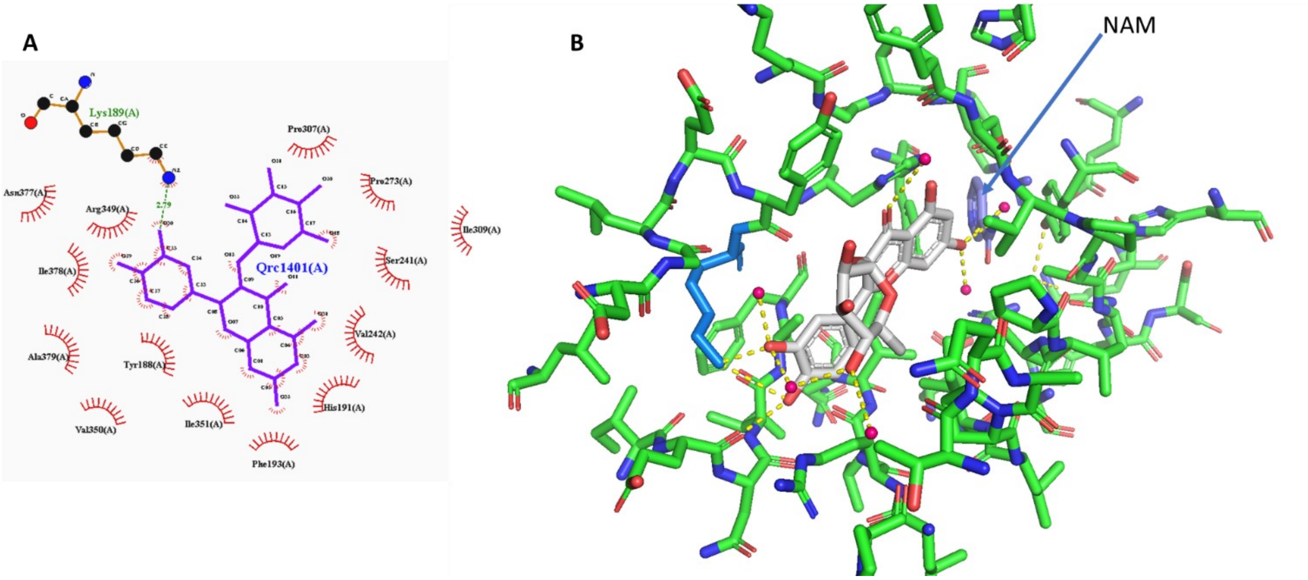

**Figure S5.** A) LigPlot* v2.2.5(15, 16) representation of rear channel residues interacting with quercitrin. B) Quercitrin (silver) showing H-bonding to Lys-189 (blue) and multiple water molecules in the rear channel

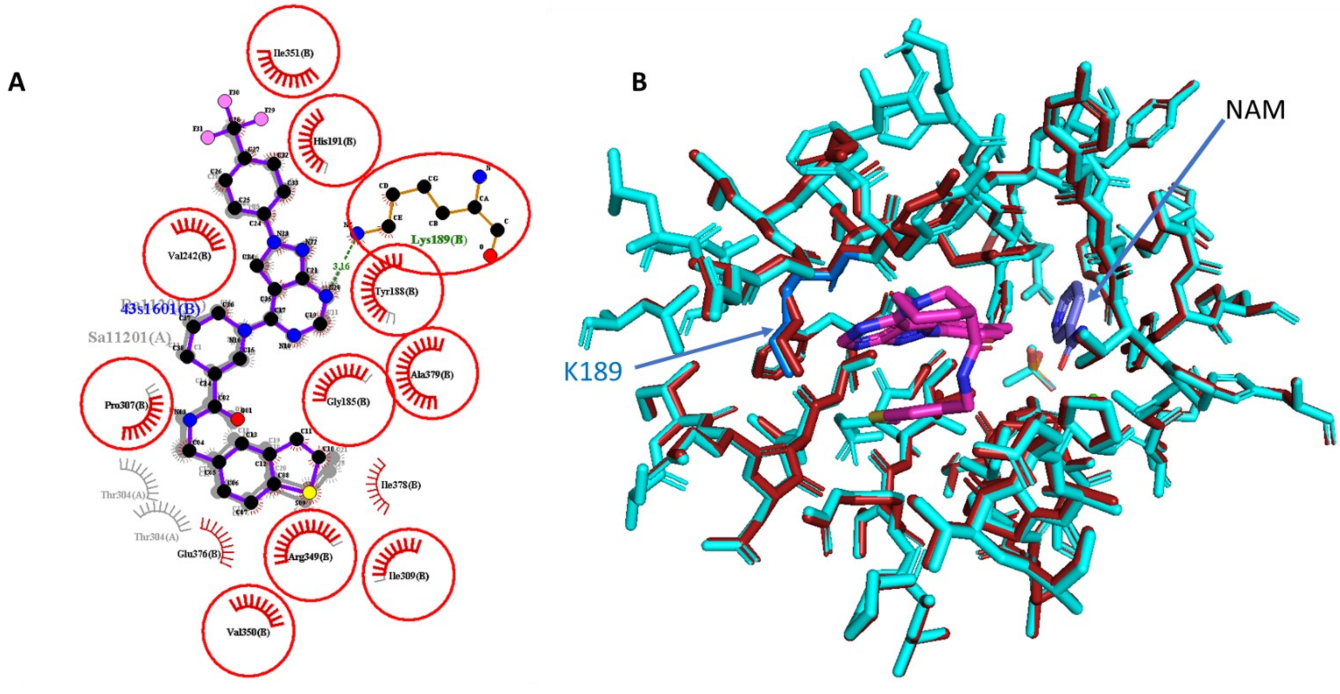

**Figure S6.** A) LigPlot* v2.2.5 superposition of all three N-PAMs bound to the rear channel showing H-bonding with Lys-189 and rear channel residues interacting with N-PAMs. B) Superposition of the co-crystal structure of NP-A1R (magenta ligand and red residues) with NAM bound to NAMPT with the structure of NAM bound without any ligand in the rear channel (cyan residues). N-PAM binding causes minimal perturbation of the amino acid positions and NAM.

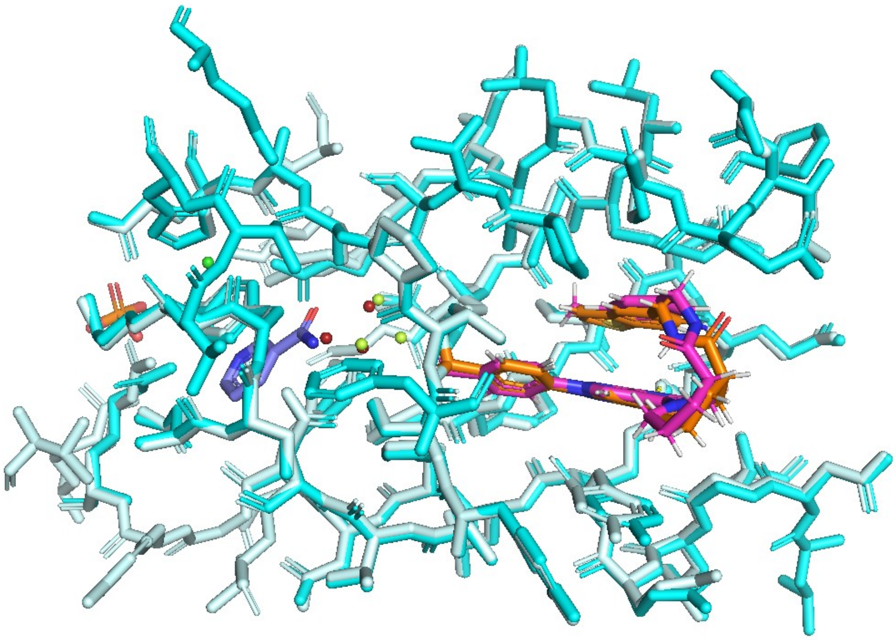

**Figure S7.** Superposition of ZN-2-43 and NP-A1S bound to the rear channel of NAMPT with NAM from the ZN-2-43 structure in blue. The three water molecules of the H-bonding network are in lime (NP-A1S) and red (ZN-2-43) and almost superimposable. A fourth water molecule is seen in the nucleobase pocket in the absence of NAM.

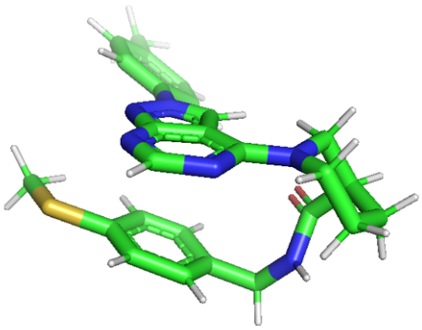

| Structures optimized at B3LYP/6-31+G*//B3LYP/6-31+G* in the gas phase | | | |
|---|---|---|---|
| | ΔE kcal/mol | ΔH kcal/mol | ΔG kcal/mol |
| NP-A1R | 0 | 0 | 0 |
| NP-A1S | 1.30 | 1.04 | 1.38 |

**Figure S8.** Structures and relative energy of calculated local energy minima, at the indicated level of theory, for N-PAM isomers. Initial coordinates were extracted from co-crystal structures .

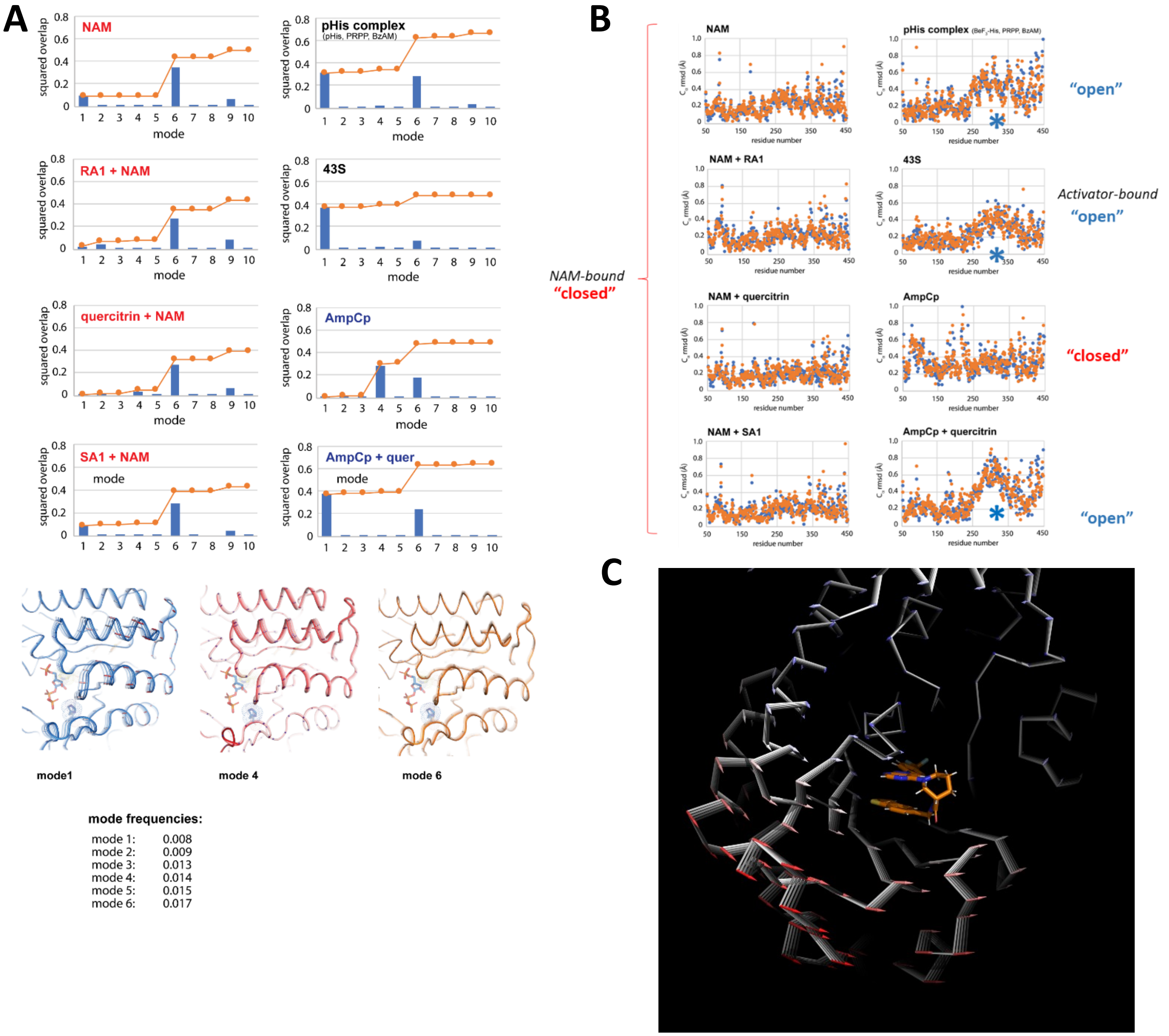

**Figure S9. A.** NMA of NAMPT crystal structure showing relative contributions of each vibrational mode and pictorial representation. **B.** Root mean deviation for Cα of each amino acid residue of NAMPT from NMA analysis of crustal structures. **C.** A movie showing Mode 1 motions is provided in a separate pptx supplemental file.

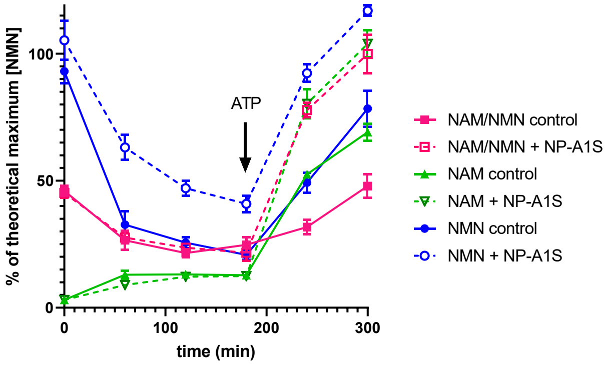

**Fig. S10.** NAMPT was incubated with various combinations of substrate, product, and N-PAM before addition of ATP. In all cases, the equilibrium position favours NAM and addition of ATP shifts the equilibrium from NAM to formation of NMN. The presence of N-PAM accelerates the turnover of NAM to NMN after addition of ATP. Time zero represents addition of either NAM (green), NMN (blue), or a combination (pink). ATP was added at 180 min. In the absence of ATP, an equilibrium position was reached favoring NAM, with addition of ATP shifting the equilibrium towards NMN. In the presence of NP-A1S, the formation of NMN was accelerated under all conditions. ATP (2 mM), PRPP 100uM, PPi 100uM, NAMPT 40nM, NAM 3uM, NMN 3uM, SBI 5uM, NP-A1S 20uM.

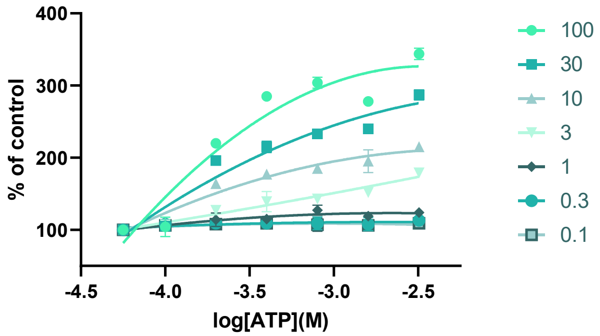

**Fig. S11.** ATP dependence of NAMPT activation by different concentrations of NAT1 (μM), measured by the orthogonal enzyme activity assay.

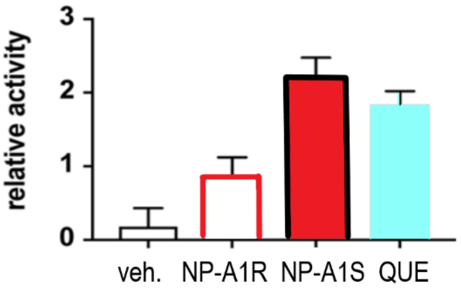

**Fig. S12.** Feedback inhibition of NAMPT activity by $NAD^+$ measured in the primary enzyme assay in the presence of $NAD^+$ (10 μM): inhibition was attenuated in the presence of N-PAMs (10 μM) and quercitrin (50 μM).

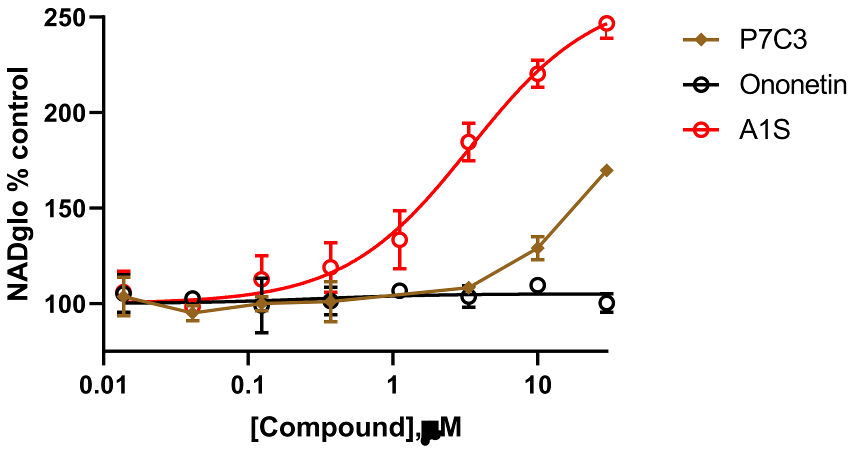

**Fig. S13.** NAD+ regulation in THP-1 cells: Concentration-response for P7C3-A20 compared to NP-A1S. The phenol ononetin was not active in THP-1 cells.

| | NAMPT complex | | | | | | |
|---|---|---|---|---|---|---|---|
| **Data Collection** | **NP-A1-R + NAM** | **NP-A1-S + NAM** | **quercitrin + NAM** | **NAM** | **quercitrin + AMPcP** | **ZN-2-43S** | **ZN-4-3** |
| PDB ID | **8DSC** | **8DSD** | **8DSE** | **8DSI** | **8DSH** | **8DTJ** | **8DSM** |
| Space Group | $P1\,2_1\,1$ | $P1\,2_1\,1$ | $P1\,2_1\,1$ | $P1\,2_1\,1$ | $P1\,2_1\,1$ | $P1\,2_1\,1$ | $P2_1\,2_1\,2_1$ |
| Cell Dimensions | | | | | | | |
| a,b,c (Å) | 60.38 106.32 82.79 | 60.51 106.69 82.80 | 60.87 106.79 82.69 | 60.82 106.63 83.03 | 60.86 107.27 83.52 | 60.50 107.48 83.40 | 87.99 94.42 246.45 |
| α,β,γ (º) | 90.00 96.45 90.00 | 90.00 96.55 90.00 | 90.00 96.61 90.00 | 90.00 96.70 90.00 | 90.00 96.78 90.00 | 90.00 96.64 90.00 | 90.00 90.00 90.00 |
| Resolution (Å) | 82.26 (1.32)* | 82.26 (1.43) | 82.14 (1.43) | 82.46 (1.43) | 82.94 (2.20) | 82.84 (2.12) | 123 (2.75) |
| Unique Observations | 245,312 (11,964) | 190,322 (7,629) | 190,913 (6,809) | 191,396 (7,807) | 52,654 (4,337) | 59,407 (3,923) | 54,145 (4,379) |
| Completeness (%) | 99.9 (99.1) | 98.9 (80.0) | 98.5 (70.8) | 99.0 (81.7) | 97.5 (92.8) | 98.5 (84.0) | 99.7 (100.0) |
| Redundancy | 3.7 (3.2) | 3.7 (1.8) | 3.5 (1.7) | 3.6 (1.7) | 3.0 (1.9) | 3.7 (1.8) | 7.4 (7.6) |
| $R_{merge}$ | 0.065 (0.655) | 0.042 (0.539) | 0.081 (0.454) | 0.055 (0.850) | 0.085 (0.427) | 0.084 (0.499) | 0.188 (1.391) |
| I/σI | 9.6 (1.8) | 15.6 (1.4) | 7.7 (1.1) | 11.8 (0.8) | 7.6 (1.7) | 9.8 (1.4) | 10.1 (1.7) |
| Source | LS-CAT ID-F | LS-CAT ID-F | LS-CAT ID-F | LS-CAT ID-F | LS-CAT ID-G | LS-CAT ID-F | LS-CAT ID-F |
| Collection date | 8/17/18 | 6/22/19 | 6/22/19 | 8/16/19 | 10/19/19 | 12/1/19 | 12/20/21 |
| Wavelength | 0.9787 | 0.9787 | 0.9787 | 0.9787 | 0.9786 | 0.9787 | 0.9787 |
| **Refinement** | | | | | | | |
| Resolution (Å) | 1.32 | 1.43 | 1.43 | 1.43 | 2.20 | 2.12 | 2.75 |
| $R_{work}$ ; $R_{free}$ (%) | 15.9; 17.5 | 16.4; 18.6 | 16.7; 18.5 | 16.1; 18.0 | 16.7; 21.2 | 21.6; 24.6 | 19.6; 23.9 |
| Number of atoms (protein/other/solvent) | 7584/ 118/ 1038 | 7544/105/977 | 7494/95/972 | 7588/37/933 | 7542/118/554 | 7473/105/408 | 14,929/229/64 |
| B-factors (Å²) (protein/other/water) | 14.3/18.6/26.9 | 18.2/25.0/30.3 | 17.1/20.0/28.3 | 19.0/23.4/31.3 | 27.7/37.9/32.5 | 36.8/43.0/40.3 | 59.1/61.2/49.1 |
| Rmsd bond lengths (Å) | 0.013 | 0.013 | 0.007 | 0.012 | 0.012 | 0.004 | 0.008 |
| Rmsd bond angles (º) | 1.50 | 1.30 | 0.88 | 1.25 | 1.22 | 0.82 | 1.06 |
| Ramachandran | | | | | | | |
| Favored | 98.39 | 98.05 | 98.06 | 98.17 | 97.86 | 98.07 | 98.71 |
| Allowed | 1.61 | 1.95 | 1.94 | 1.83 | 2.14 | 1.93 | 1.29 |
| Forbidden | 0.00 | 0.00 | 0.00 | 0.00 | 0.00 | 0.00 | 0.00 |
| Rotamer outliers (%) | 0.24 | 0.24 | 0 | 0.48 | 0.12 | 0.12 | 1.00 |
| Molecules in ASU | 2 | 2 | 2 | 2 | 2 | 2 | 4 |
| **Programs Used** | | | | | | | |
| Processing | HKL2000 | XDS | XDS | XDS | DIALS | XDS | XDS |
| Scaling | HKL2000 | XDS | XDS | XDS | Aimless | XDS | XDS |
| Phasing | Molrep | Molrep | Molrep | Molrep | Molrep | Molrep | Molrep |
| Phasing Model | 2E5D | 2E5D | 2E5D | 2E5D | 2E5D | 2E5D | 2E5D |
| Manual Build | Coot | Coot | Coot | Coot | Coot | Coot | Coot |
| Refinement | Phenix | Phenix | Phenix | Phenix | Phenix | Phenix | Phenix |

* values in parentheses are for the highest resolution shell

### References

1. Studier FW (2005) Protein production by auto-induction in high density shaking cultures. *Protein Expr Purif* 41(1):207-234.
2. Burgos ES & Schramm VL (2008) Weak coupling of ATP hydrolysis to the chemical equilibrium of human nicotinamide phosphoribosyltransferase. *Biochemistry* 47(42):11086-11096.
3. Kalgutkar AS*, et al.* (2005) A comprehensive listing of bioactivation pathways of organic functional groups. *Curr Drug Metab* 6(3):161-225.
4. Erve JC (2006) Chemical toxicology: reactive intermediates and their role in pharmacology and toxicology. *Expert Opin Drug Metab Toxicol* 2(6):923-946.
5. Zhang RY*, et al.* (2011) A fluorometric assay for high-throughput screening targeting nicotinamide phosphoribosyltransferase. *Anal Biochem* 412(1):18-25.
6. Kabsch W (2010) Xds. *Acta Crystallogr D Biol Crystallogr* 66(Pt 2):125-132.
7. Winter G (2010) xia2: an expert system for macromolecular crystallography data reduction. *J Appl Cryst* 43(1):186-190.
8. Winter G*, et al.* (2018) DIALS: implementation and evaluation of a new integration package. *Acta Crystallogr D Struct Biol* 74(Pt 2):85-97.
9. Vagin A & Teplyakov A (2010) Molecular replacement with MOLREP. *Acta Crystallogr D Biol Crystallogr* 66(Pt 1):22-25.
10. Adams PD*, et al.* (2010) PHENIX: a comprehensive Python-based system for macromolecular structure solution. *Acta Crystallogr D Biol Crystallogr* 66(Pt 2):213-221.
11. Afonine PV*, et al.* (2012) Towards automated crystallographic structure refinement with phenix.refine. *Acta Crystallogr D Biol Crystallogr* 68(Pt 4):352-367.
12. Emsley P, Lohkamp B, Scott WG, & Cowtan K (2010) Features and development of Coot. *Acta Crystallogr D Biol Crystallogr* 66(Pt 4):486-501.
13. Pearce NM, Krojer T, & von Delft F (2017) Proper modelling of ligand binding requires an ensemble of bound and unbound states. *Acta Crystallogr D Struct Biol* 73(Pt 3):256-266.
14. Grant BJ, Rodrigues AP, ElSawy KM, McCammon JA, & Caves LS (2006) Bio3d: an R package for the comparative analysis of protein structures. *Bioinformatics* 22(21):2695-2696.
15. Laskowski RA & Swindells MB (2011) LigPlot+: multiple ligand-protein interaction diagrams for drug discovery. *J Chem Inf Model* 51(10):2778-2786.
16. Wallace AC, Laskowski RA, & Thornton JM (1995) LIGPLOT: a program to generate schematic diagrams of protein-ligand interactions. *Protein Engineering* 8(2):127-134.
